# Isotopic signatures induced by upwelling reveal regional fish populations in Lake Tanganyika

**DOI:** 10.1101/2021.10.05.463178

**Authors:** Benedikt Ehrenfels, Julian Junker, Demmy Namutebi, Cameron M. Callbeck, Christian Dinkel, Anthony Kalangali, Ismael A. Kimirei, Athanasio S. Mbonde, Julieth B. Mosille, Emmanuel A. Sweke, Carsten J. Schubert, Ole Seehausen, Catherine E. Wagner, Bernhard Wehrli

**Author notes:** Corresponding author: Benedikt Ehrenfels. shared first authorship (Benedikt Ehrenfels, Julian Junker).

## Abstract

Lake Tanganyika’s pelagic fish sustain the second largest inland fishery in Africa and are under pressure from heavy fishing and global warming related increases in stratification. Only little is known about whether basin-scale hydrodynamics – including a more stratified north and an upwelling-driven south – induce ecological and genetic differences among populations of highly mobile, pelagic fish inhabiting these different areas. Here, we examine whether the basin-scale dynamics leave distinct isotopic imprints in the pelagic fish of Lake Tanganyika, which may reveal differences in habitat, diet, or lipid content. We conducted two lake-wide campaigns during different seasons and collected physical, nutrient, chlorophyll, phytoplankton and zooplankton data. Additionally, we analyzed the pelagic fish – the clupeids *Stolothrissa tanganicae*, *Limnothrissa miodon* and four *Lates* species – for their isotopic and elemental carbon (C) and nitrogen (N) compositions. The δ^13^C values were significantly higher in the productive south after the upwelling/mixing period across all trophic levels, implying that the fish have regional foraging grounds, and thus record these latitudinal isotope gradients. By combining our isotope data with genetics, we demonstrate that the fish form regional populations on a seasonal to multiannual time scale. Based on δ^15^N and C:N ratios, we found no strong evidence for varying diets or lipid contents between those regional populations.

Additional analyses revealed that isotopic variations between specimens from the same location are not linked to genetic differences. We suggest that the development of basinscale ecological differences in response to the prevailing hydrodynamic regimes may be inhibited by lake-wide gene flow on the long term. Our findings show that the pelagic fish species are genetically adapted to the whole lake, but they form regional populations on short time scales. This implies that sustainable management strategies may adopt basin-scale fishing quotas.

## 1 Introduction

Lake Tanganyika is by volume the second largest freshwater lake in the world, and its pelagic fish community sustains the second largest inland fishery in Africa (1), providing important employment opportunities and animal protein for millions of people in the riparian communities (2, 3). The pelagic food web in Lake Tanganyika is composed of a copepod-dominated zooplankton assemblage, a phyto- and zooplankton grazer community consisting of two endemic sardine species (*Stolothrissa tanganicae* and *Limnothrissa miodon*), and a predator assemblage comprising of four endemic latid species (genus *Lates*), of which *Lates stappersii* is the most common (4). Today, the sardines and *Lates stappersii* account for 95 % of the pelagic fish catch in Lake Tanganyika (5).

The pelagic fish stocks suffer from heavy fishing (2, 6) and from a long term decline that was attributed to climate change (7–9). The increased warming of the surface waters caused by climate change leads to steep temperature gradients in the water column. These gradients build physical barriers to vertical mixing, thereby limiting the transfer of nutrients to surface waters where light is available to drive primary productivity (7,8,10– 13).

Although assessing potential long-term changes in the ecology of the pelagic fish is impaired by data scarcity, Lake Tanganyika’s limnological cycle offers the opportunity to study the impact of varying levels of stratification on a basin-scale. This annual cycle is driven by climatic differences between the north and south and is characterized by four stages (Fig. 1a-d; 14,15): (i) In the warm rainy season (November-March), stagnant and highly stratified waters lead to an overall nutrient-depleted epilimnion (Fig. 1 a), and internal waves only cause local nutrient injections into the surface waters (16). (ii) In March-May, the southeast trade winds initiate the lake circulation in the upper water column, resulting in strong nutrient upwelling in the southern basin (Fig. 1 b). (iii) The upwelling in the south transforms into a convective mixing of the upper ∼150 m. The sinking, cool surface waters in the south reverse the lake circulation by initiating a northward current between 50-100 m and a surface counter current, further weakening thermal stratification across the lake (Fig. 1 c). (iv) The trade winds cease in October, slowing down the circulation, while the water column re-stratifies lake-wide. A weaker, secondary upwelling leads to a nutrient pulse at the northern end of the lake (Fig. 1 d).

**Fig. 1:**
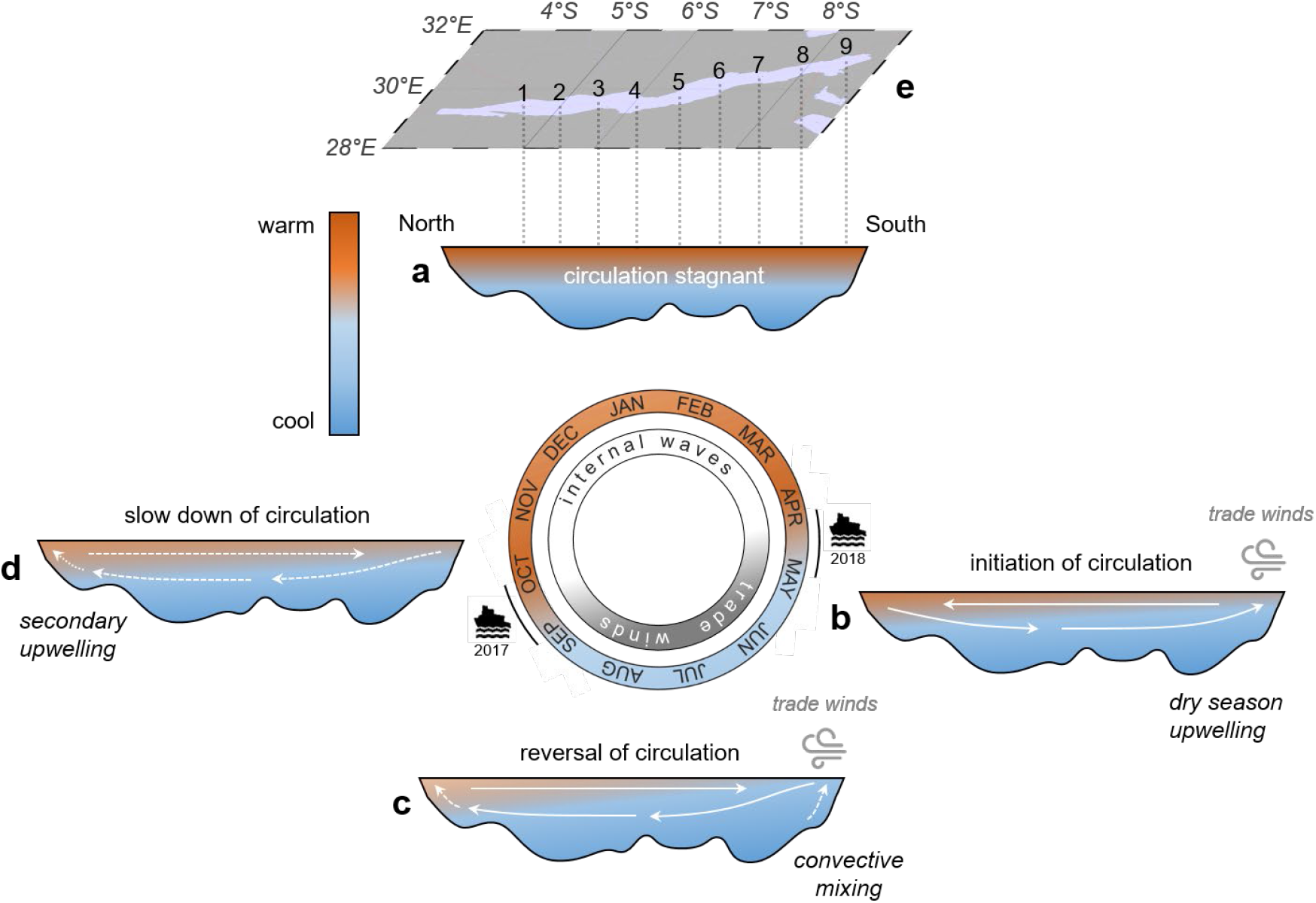
The limnological cycle of Lake Tanganyika with its four major phases according to Plisnier et al. (14) and Verburg et al. (15). (**a**) Stagnant, highly stratified waters during the warm rainy season (November-March) only support low nutrient availability. (**b**) The onset of the cool dry winds in March-May initiates the upwelling in the South leading to high nutrient fluxes in this region. (**c**) The lake circulation reverses during the dry season (May-September). Water column stratification is low and the nutrient availability high across the lake, with a maximum in the convective mixing area in the south. (**d**) The trade winds cease in October slowing down the lake circulation, while the water column re-stratifies. A weaker secondary upwelling leads to a nutrient pulse at the northern end of the lake. During the dry season, wind-driven upwelling and mixing are the dominant driving force behind nutrient injections into the euphotic zone, whereas internal waves are particularly important in the rainy season. The color gradient indicates the level of thermal stratification. Note that this latitudinal cross-section is not to scale and that the outlined mechanism primarily affects the upper water column (<200 m). Our two sampling campaigns were timed at the seasonal transitions in September/October and April/May to compare the effects of the preceding dry and rainy seasons. (**e**) The map shows the nine stations for water column and plankton sampling. Fish samples representing the pelagic catch were collected from the respective coastal villages/towns.

The limnological cycle of Lake Tanganyika leads to seasonal and regional gradients in nutrient availability, causing overall higher primary productivity in the dry season compared to the rainy season and higher primary productivity in the south than in the north of the lake (17–21). High densities of both zooplankton and the sardines *Stolothrissa* and *Limnothrissa* are coupled to the phytoplankton blooms (22–24), which also seem to benefit sardine spawning and recruitment (25). The north-south variability furthermore affects the zooplankton community composition: shrimps and calanoid copepods prevail in the south (26–28), whereas cyclopoid copepods and jellyfish dominate in the north (24,29,30). Differences in the zooplankton community may in turn influence predatory fish. Mannini et al. (31) found that the diet of *Lates stappersii* in the north is heterogeneous and consists of copepods, shrimps, and sardines, whereas *Lates stappersii* in the south feed mainly on shrimps.

The spatial variability in Lake Tanganyika’s pelagic habitat, driven by the mixing regime, could additionally impact the life cycle (e.g. spawning phenology, developmental timing and recruitment success) of the pelagic fish species (23,25,32,33). The spatial variation in the pelagic environment might generate different fitness optima and drive divergent adaptation of pelagic fish populations between north and south, if horizontal migration remains limited. However, recent genetic studies of the sardines (34, 35) and the four *Lates* species (36) did not find evidence for genetic population differentiation along the north-south gradient, suggesting that gene flow may overcome any possible effects of divergent natural selection between the basins. Nonetheless, the fish populations may still respond to differences in physicochemical conditions and food supply by evolving ecological adaptations to the regional environments. Such responses might involve variations in food web interactions (37) or lipid storage for bridging lean periods (38, 39). However, possible effects of the seasonality and spatial variation in the physical mixing regime of Lake Tanganyika on the distribution and ecology of its pelagic fish have not been studied.

Characterizing seasonal and spatial gradients in the carbon (C) and nitrogen (N) elemental and isotopic composition of Lake Tanganyika’s food web may provide insight to the migration distances, diets, and lipid contents of the fish species. Indeed, an isotopic study of fish and their surrounding food web along a geographical gradient can reveal regional population isolation if environmental differences among sites translate into divergent isotopic signatures of regional or local fish populations (40). In this isotopic framework, the ^13^C/^12^C ratio or δ^13^C increases only little from one trophic level to the next and therefore reflects the source of primary production (41–43). Differences in primary productivity can alter the δ^13^C of particulate organic matter (POM): high primary productivity results typically in high values of δ^13^C, due to the ongoing depletion of the of the DIC pool and decreasing discrimination against ^13^C by phytoplankton (44–49).

Changes in δ^13^C at the base of the food web can be tracked across trophic levels from plankton to fish in lake ecosystems (50). In Lake Tanganyika, O’Reilly et al. (7) and Verburg (12) used the δ^13^C-POM to reconstruct historical changes in primary productivity. In addition, previous δ^13^C analyses in the northern basin have reported higher δ^13^C-POM values in the productive dry season compared to the rainy season (51–53). The ratio of ^15^N/^14^N, or δ^15^N, provides insight into the trophic position of an organism, because it increases significantly with each trophic level. This successive enrichment allows estimating an organism’s trophic position in the food web and is used in ecology to describe prey and predator relationships (41–43). Lastly, the elemental C:N ratio was often used to infer the lipid content of fish muscle tissue, with higher C:N denoting higher lipid contents (54, 55).

In this study, we explore the latitudinal and seasonal patterns of δ^13^C and δ^15^N in the pelagic food web of Lake Tanganyika in the context of the lake’s limnological variability. During two lake-wide field campaigns in the final phases of the dry and the rainy seasons, we measured the C and N isotopic and elemental compositions of the major pelagic food web members (POM, zooplankton, the bivalve *Pleiodon spekii*, fish). These samples were collected in concert with limnological data including the physical properties of the water column, oxygen and nutrient concentrations, chlorophyll, as well as the phyto- and zooplankton community and abundance (56). Using the extensive data sets from those two contrasting time points, we first tested to which extend the regional and seasonal patterns in primary productivity induce systematic differences in the isotopic signatures of plankton, and then tracked the isotopic signals and C:N ratios through the food web to the pelagic fish. The results allowed us to assess the extent of regional isolation and ecological differentiation of the pelagic fish populations. Finally, we tested whether existing genetic differences (35, 36) were linked to dietary differences in the six major pelagic fish species.

## 2 Materials & Methods

### 2.1 Study site and sampling

Our two Lake Tanganyika sampling campaigns, spanning two different hydrological conditions across a north-south transect of ∼500 km, were conducted at the end of the dry season (28 September - 8 October 2017) and the end of the following rainy season (27 April - 7 May 2018). Water column and plankton characteristics were sampled during two cruises on *M/V Maman Benita* (56, 57). At the end of the dry season, we collected fish samples as described in Junker et al. (2020) at station 1, station 2, station 5, station 7 and station 9 during a land-based excursion prior to the cruise (17-24 September 2017), whereas all nine landing sites, corresponding to our nine pelagic sampling stations, were sampled during the cruise at the end of the rainy season (Fig. 1 e). In addition, we took fish samples in Kigoma in July 2017 (35).

### 2.2 Physical and chemical parameters

We measured temperature, dissolved oxygen, photosynthetically active radiation, and in-situ chlorophyll fluorescence via CTD profiling (Sea-Bird SBE 19plus) at each station. From these stations, we also collected water with large Niskin bottles (20-30 L) at 5-25 m depth intervals down to 250 m depth. Water column stratification was expressed as buoyancy frequency (N^2^) and Schmidt stability (Sc). We interpreted clear peaks in N^2^ as thermoclines, whereby the N^2^ value at the peak provides a measure of steepness of the thermocline. In addition, we calculated the Sc over 1 m^2^ between 50 and 100 m for each station using the R package ‘rLakeAnalyzer’ (58). This depth interval extends from the typical location of the nitrate peak to the bottom of the euphotic zone (56,59,60). For a more detailed description of the thermal structure of the water column see Ehrenfels et al. (56).

Water samples to measure nutrients (phosphate, ammonium, nitrate, and nitrite) were taken directly from the Niskin bottles, filtered sterile through 0.2 µm filters and processed on-board following standard methods (61–63). On average, the detection limits were 0.22, 0.34, 0.20, and 0.03 µM for phosphate, ammonium, nitrate, and nitrite, respectively.

Water samples to measure dissolved inorganic carbon (DIC) were collected in 12 mL exetainers directly from the Niskin bottles and filtered sterile (0.2 µm). Samples were stored at room temperatures and shipped to Switzerland. At Eawag Kastanienbaum, the DIC concentrations were measured by high temperature combustion catalytic oxidation using a Shimadzu TOC-L Analyzer (Shimadzu TOC-VCPH/CPN). A 2 mL aliquot of the DIC sample was used to quantify the isotopic fractionation of δ^13^C-DIC. The aliquot subsample was transferred to a new 12 mL exetainer, where it was Helium purged for 2 minutes. The sample was then capped and 50 μL orthophosphorous acid (85 %) was added. The samples were mixed and stored for ∼15 h at room temperature for equilibration prior to analysis by GC-IRMS (Isoprime). Sample δ^13^C-DIC was adjusted to the standard *Carrara marmor* (ETH Zurich).

### 2.3 CO2 fixation rates

Carbon fixation incubations and rate calculations were done as described in Schunck et al. (64) and Callbeck et al. (65). Briefly, samples were carefully filled from the Niskin into 4.5 L polycarbonate bottles capped with polypropylene membranes. Per sampled depth, we filled off triplicate bottles, including one control (no added label) and duplicate treatments (with amended ^13^C-HCO_3-_). We added 4.5 mL of ^13^C-bicarbonate solution (1 g ^13^C-bicarbonate in 50 ml water; Sigma Aldrich) to each of the treatment bottles. The label was mixed in the treatment bottles for ∼30 min under shaking. Thereafter, a 12 mL subsample was taken for quantifying the labelling percent (mean 2.8 %). The resulting headspace was re-filled with water from the same depth, and bottles were then incubated headspace-free in 60 L incubators covered with shaded light filters (LEE Filters) mimicking the in-situ irradiance and light spectrum. After 24 h, the samples were filtered on pre-combusted GF/F filters (Whatman). The filters were oven-dried (60° C for 48 h) and stored at ambient temperatures. Filter samples were shipped to Switzerland and further processed as described in 2.7. Due to the small difference between the in-situ and incubation temperatures (<5° C), the derived CO_2_ fixation rates were not adjusted for temperature.

### 2.4 Chlorophyll, phytoplankton and particulate matter

We measured the chlorophyll-*a* concentrations according to Wasmund, Topp & Schories (66). Briefly, 2-4 L of lake water were filtered through 47 mm glass fibre filters (GF55, Hahnemühle), which were directly transferred to 15 mL plastic tubes. Five mL ethanol (>90 %) were added to the samples, followed by 10 min cold ultrasonification. The samples were stored at 5 °C overnight and sterile-filtered (0.2 µm) the following morning. The extracts were measured on-board with a fluorometer (Turner Trilogy) and calibrated against a chlorophyll-*a* standard (Lot# BCBS3622S, Sigma-Aldrich). Samples and standards were always handled and processed in the dark. In-situ chlorophyll fluorescence was calibrated against extracted chlorophyll-*a* samples and then used to calculate depth-integrated chlorophyll-*a* stocks (0-125 m).

For estimating the phytoplankton abundances, 4-10 L of water were concentrated to 20 mL using a 10 μm plankton net and fixed with alkaline Lugol solution. At TAFIRI Kigoma, phytoplankton cells were counted from 2 mL subsamples by inverted microscopy (at ×400 magnification). For particulate organic matter (POM), 2-4 L lake water was filtered through precombusted GF/F filters (nominal pore size 0.7 µm; Whatman).

### 2.5 Zooplankton and Pleiodon spekii

Zooplankton was collected with vertical net hauls across the oxygenated water column (0-150 m) at each pelagic station. We sampled different size fractions of the zooplankton community using three different nets. For smaller zooplankton, we used 25 and 95 µm nets with 0.03 and 0.02 m^2^ mouth openings, respectively. A 250 µm net with a 0.28 m^2^ mouth opening was used for larger, fast swimming species. We preserved all zooplankton collected from the first haul in ethanol for taxonomic zooplankton community assessment, while the individuals from the second haul were designated for stable isotope analysis (only for samples from the 95 and 250 μm nets). At TAFIRI Kigoma, we analyzed the zooplankton community composition of the ethanol-preserved samples by compound microscopy (Leica Wild M3B) at x200 magnification. Additionally, we picked living individuals of the long-lived, filter-feeding bivalve *Pleiodon spekii* at near-shore habitats in water depths of 1.5-6 m by snorkelling. Bivalves were first euthanized with an overdose of MS222. Then we sampled the foot using clean scalpels and forceps and removed the mucous with tissues and deionized water.

### 2.6 Fish

At on-shore landing sites adjacent to our sampling stations, we obtained fish specimens from fishermen, which usually fish within a 20 km radius from their landing sites. We collected *Stolothrissa tanganicae*, *Limnothrissa miodon*, *Lates stappersii*, *Lates microlepis*, *Lates mariae*, and *Lates angustifrons* and processed them according to the standard protocol described in Junker et al. (35). For stable isotope analysis, we sampled the dorsal muscle using clean scalpels and forceps and removed the skin.

### 2.7 Isotopic and elemental analysis of solids

All solid isotope samples (POM, zooplankton, *P. spekii*, and fish) were oven-dried at ∼60°C for at least 24 h after collection and then packed in aluminium foil or small sample tubes. Dried samples were stored at room temperature and shipped to Switzerland. At Eawag Kastanienbaum, we fumed the POM samples for 48 h under HCl atmosphere to remove inorganic carbon. Fish and *P. spekii* samples were ground to fine powder using a Qiagen Tissuelyzer II. We measured the C and N elemental and isotopic compositions with an EA-IRMS (vario PYRO cube, Elementar coupled with an IsoPrime IRMS, GV Instruments). *Acetanilide #1* (Indiana University, CAS # 103-84-4) was used as an internal standard. The isotopic ratios of the samples are reported in the delta notation VPDB for carbon and air for nitrogen. Standard and sample reproducibility was generally better than 0.2 ‰ for δ^13^C and 0.5 ‰ for δ^15^N and highest for fish tissue (0.1 ‰ for δ^13^C and 0.2 ‰ for δ^15^N).

### 2.8 Lipid content of fish muscle tissue

For a subsample of *Stolothrissa* individuals, which exhibited the largest range in C:N ratios, we measured the total lipid content to test whether a high C:N ratio effectively translates to a higher amount of lipids in fish tissue. Total lipid content was determined gravimetrically following Folch, Lees & Sloane-Stanley (67). In brief, ∼1 mg of dried fish muscle powder was weighed into a pre-combusted glass vial, and 1 mL of 2:1 (vol:vol) dichloromethane:methanol solution was added. The sample was then ultrasonicated for 10 min. The supernatant was transferred to another pre-combusted, pre-weighed glass vial and evaporated in a heat block. The entire procedure was repeated two more times, and the resulting dry lipid mass weighed to the nearest 0.001 mg.

### 2.9 Data analysis

For calculating the depth-integrated isotopic values of POM, we normalized each sample for the phytoplankton abundance at the respective depth. We corrected the δ^13^C of non-lipid-extracted animal tissue for its lipid content according to Post et al. (54). We estimated lipid content in fish tissue according to the theoretical model from the same study. For comparing the isotopic composition and the C:N ratios of fish populations between different regions, we selected individuals from stations where samples were available from both campaigns (north: stations 1 and 2; south: stations 7 and 9). From this subset, we additionally selected the individuals from the 50 mm size range with the highest overlap across regions and sampling campaigns for each species to minimize size-specific effects (Figs. S1; S2; 62). The chosen size ranges were 40-90 mm for *Stolothrissa*, 75-125 mm for *Limnothrissa*, and 200-250 mm for *Lates stappersii*. For the same analyses and reasons, we selected *P. spekii* from the exact same sites, whereas for zooplankton we used stations 1-3 (north) and 7-9 (south), because zooplankton samples were available from all stations, but we had only one sample per station. The Bayesian ellipses in the isotopic space were calculated using the R package SIBER (69).

## 3. Results

### 3.1 Biogeochemistry and hydrodynamic conditions in Lake Tanganyika during sampling

We observed strong differences in water column stratification and biogeochemistry in Lake Tanganyika between the two sampling campaigns and among basins (Fig. 2). Our sampling campaign at the end of the dry season (September/October) exhibited southerly trade winds that are characteristic for the wind-driven upwelling and mixing period (Fig. 1 c, d). By contrast, the winds were mostly calm and it was rainy at the end of the rainy season (April), which is typical for this time (Fig. 1 b). Compared to Apr/May, we find significantly lower Sc and N^2^ values as well as lower surface temperatures (Δ∼0.5 °C) in Sep/Oct, indicating that the lake was less stratified during that period (Fig. 2 a-d; Sc: 0.4-0.7 versus 0.5-1.1 kJ m^-2^; N^2^: 1.5-2.7 versus 2.4-5.0 · 10^-4^ s^-2^; Mann-Whitney-U Tests, *p* < 0.05). However, during both campaigns Sc values of the 50-100 m interval decreased towards the south, indicating some degree of upwelling/mixing (Fig. 2 a, b). At the end of the rainy season, the southern stations were sampled in May during the onset of the trade winds. The thermocline was not as heavily uplifted in the southern basin in Apr/May compared to Sep/Oct, when no thermocline had formed at station 9 (Fig. 2 c, d). This is a good indicator that wind-driven upwelling/mixing was not as pronounced in Apr/May.

**Fig. 2:**
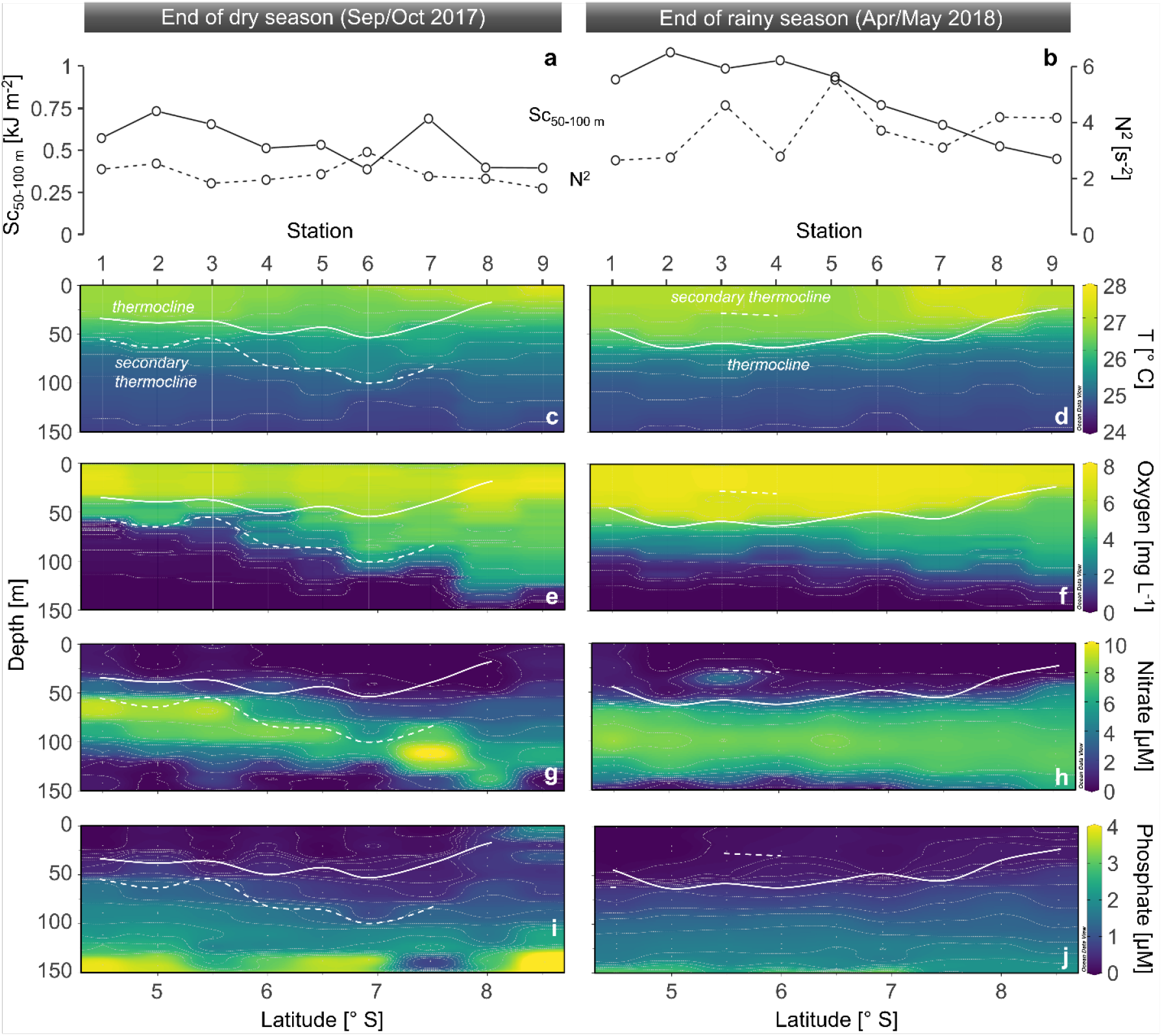
Physical and chemical properties of Lake Tanganyika along our north-south transects (from station 1-9) at the end of the dry season (left) and the end of the rainy season (right). (**a**,**b**) Schmidt stability (Sc) of the 50-100 m depth interval and buoyancy frequency of the primary thermocline (N2). Distribution of (**c**,**d**) temperature (T), (**e**,**f**) dissolved oxygen, (**g**,**h**) nitrate, and (**i**,**j**) phosphate. The solid white line depicts the thermocline, whereas the dashed white line represents less pronounced secondary thermoclines. No clear thermocline had formed at station 9 at the end of the dry season. Samples are indicated by vertical lines (continuous profiles) or points (discrete samples).

The oxygen distribution closely followed the thermal structure of the lake (Fig. 2 e, f). In Sep/Oct, the more stratified water column in the northern basin exhibited a shallower oxycline (50-70 m at stations 1-3), defined here as a sharp drop in oxygen concentrations, compared to the wind-driven mixing in the south, which introduced more oxygen to deeper layers of the water column (up to 5 mg L^-1^ at 113 m at station 9; Fig. 2 e). In Apr/May, the oxycline was lying relatively shallow across the full length of the lake from 70-120 m; oxygen concentrations were roughly 2-fold lower in the deep waters of the southern basin compared to the well-mixed conditions in Sep/Oct (∼2.3 mg L^-1^ at 113 m at station 9; Fig. 2 f).

We also observed strong latitudinal changes in the nutrient distribution in Sep/Oct. Moving from north to south, the position of the nitrate maximum in the water column deepened from 67 m to 137 m, paralleling variation of the vertical phosphate gradients (Fig. 2 g, i). The surface waters in the southern basin were also associated with generally higher nitrate and especially phosphate concentrations. For instance, at some stations in the south, surface water nitrate and phosphate concentrations reached up to 2.3 µM, whereas they were <0.5 µM at most other stations. By contrast, in Apr/May, nitrate and phosphate were more uniformly distributed across the north-south transect, and the nitrate maxima were positioned at approximately 100 m across the lake (Fig. 2 h, j). A local nitrate maximum (4.7 µM) at 40 m at station 3 in Apr/May may have been caused by an overlying cyanobacterial bloom (56). The upward tilting thermocline in the south enhanced the nutrient transport to the productive surface waters during both campaigns. However, the tilting of the thermocline as well as surface nitrate and phosphate concentrations in the south, were lower in Apr/May than in Sep/Oct (Fig. 2 c, d, g, h, i, j).

### 3.2 Concentration and isotopic composition of DIC and CO2 fixation rates in April/May

In Apr/May, we determined the DIC concentration (stations 2, 5 and 7) and C isotopic composition (stations 1, 3 and 8) in the north and the south. Like other biogeochemical parameters, neither the concentration, nor the δ^13^C of DIC showed strong latitudinal trends in Apr/May (Fig. 3), but both do exhibit pronounced vertical gradients. The DIC concentration varied between 70 and 72 mg C L^-1^ in the productive upper 50 m, followed by a sharp increase to about 74 mg C L^-1^ that flattened out with increasing depth.

**Fig. 3:**
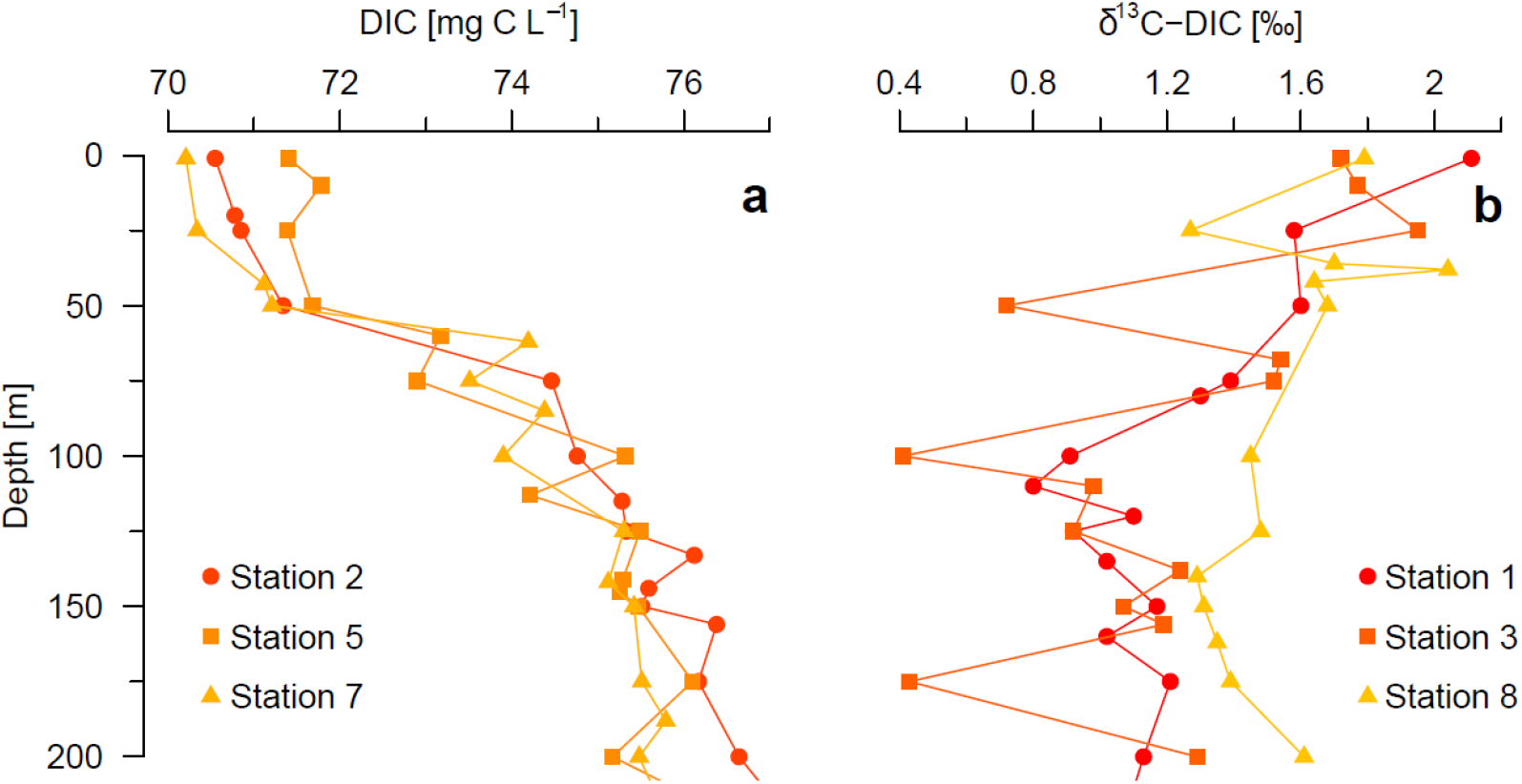
Distribution and isotopic composition of dissolved inorganic carbon (DIC) at the end of the rainy season (Apr/May 2018) in Lake Tanganyika. (**a**) DIC concentration profiles from stations 2, 5, and 7. (**b**) δ^13^C-DIC profiles from stations 1, 3, and 8.

Concurrently, the δ^13^C-DIC values reached their maximum near the surface (2.1 ‰), decreased to 0.8 ‰ at the boundary between the metalimnion and the upper hypolimnion (100-150 m) and showed a further trend to slightly heavier values at greater depth.

In addition, we performed CO_2_ fixation experiments at station 2 in the north and station 7 in the south (Fig. 4). At both stations, CO_2_ fixation rates were highest in the euphotic zone. CO_2_ fixation rates in the top 50 m were higher at station 7 in the upwelling-driven south (0.58-1.60 µM C d^-1^), compared to station 2 in the permanently stratified north basin (0.12-0.48 µM C d^-1^).

**Fig. 4:**
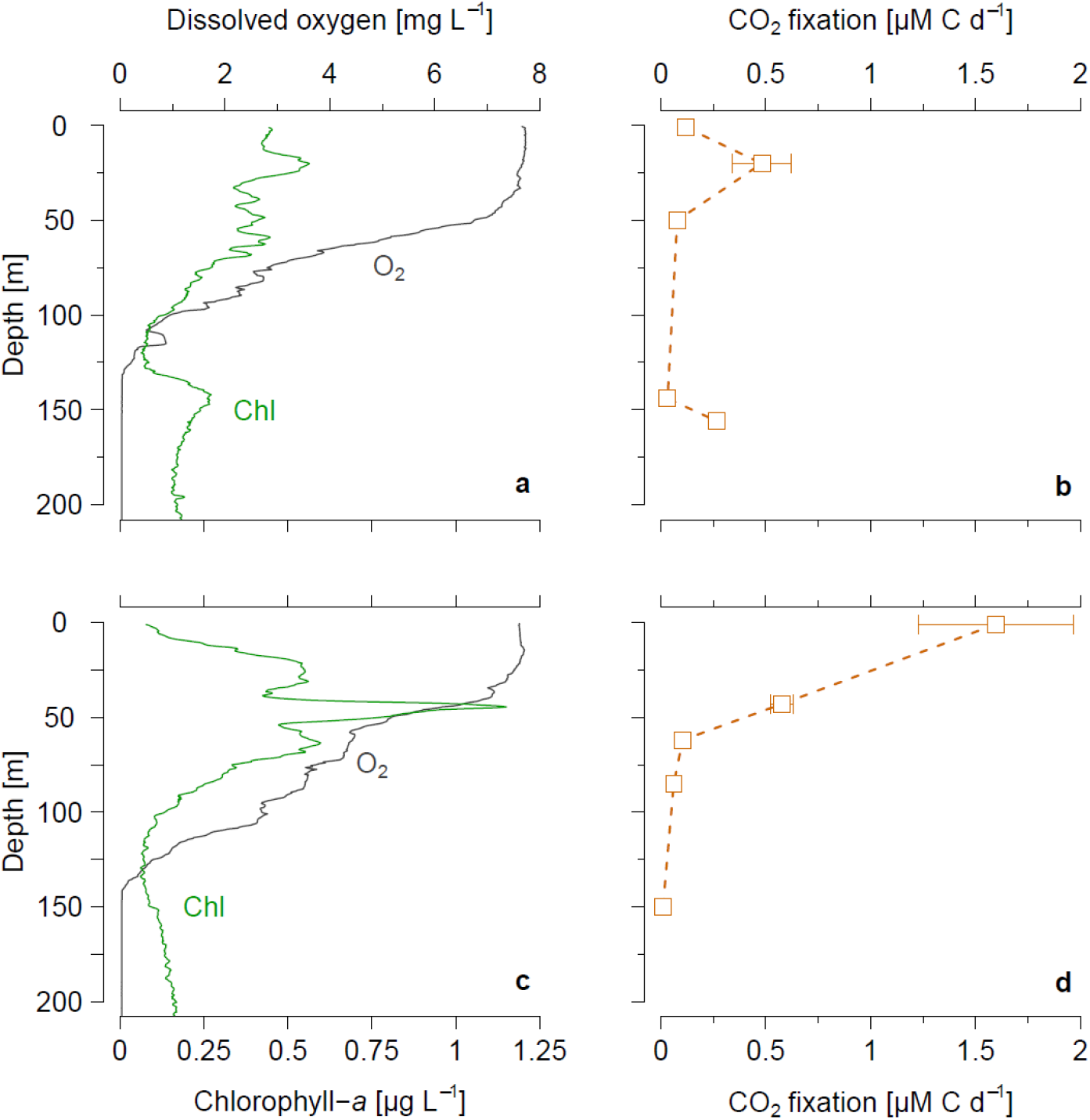
Experimentally determined CO_2_ fixation rates relative to the oxygen and chlorophyll-*a* distributions. (**a**,**c**) Oxygen and in-situ chlorophyll-*a* as well as (**b**,**d**) CO_2_ fixation rates from stations 2 (**a**,**b**) and 7 (**c**,**d**) in Apr/May.

### 3.3 Particulate matter, phytoplankton, and zooplankton

The abundance of medium-to large-celled phytoplankton (>10 µm) decreased significantly from north to south by 2.6 and 4.4 · 10^8^ ind. m^-2^ per degree latitude (linear regression, *p* < 0.05) in Sep/Oct and Apr/May, respectively (Fig. 5 a, b). Chlorophyll-*a* was slightly higher in Sep/Oct compared to Apr/May (on average 46 versus 41 mg chl-*a* m^-2^), and generally showed the highest values in the south during both campaigns (Fig. 5 a, b).

**Fig. 5:**
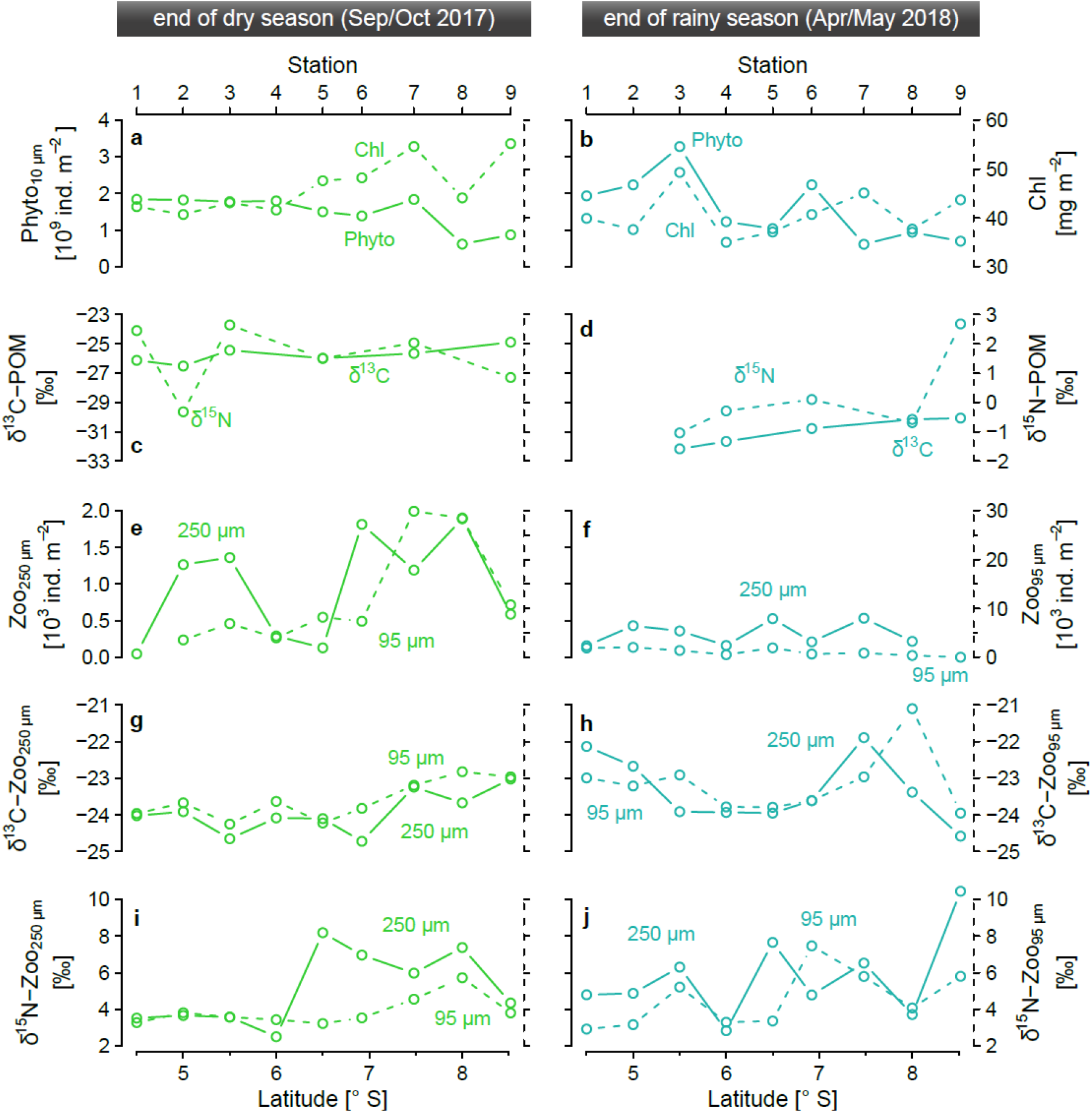
Different plankton parameters sampled across the epi- and metalimnion (0-125 m) along the north-south transects at (left) the end of the rainy season and (right) the end of the dry season. (**a**,**b**) Depth-integrated phytoplankton abundances (>10 μm) and chlorophyll-*a* stocks. (**c**,**d**) Depth-integrated and phytoplankton abundance-weighed δ^13^C and δ^15^N of POM. (**e,f**) Depth-integrated abundances of the 95 μm and 250 μm size fractions of the zooplankton community. (**g,h**) δ^13^C and (**i,j**) δ15N of the 95 μm and 250 μm size fractions of the zooplankton community

We observed strong differences in δ^13^C-POM between the two campaigns (Fig. 5 c, d). In Sep/Oct, the values of δ^13^C-POM varied between -26.5 and -24.9 ‰, whereas they spanned from -32.3 to -30.1 ‰ in Apr/May. During both seasons, we observed an increase from north to south of ∼0.3 ‰ (linear regression, *p* < 0.1) and ∼0.7 ‰ (linear regression, *p* < 0.01) per degree latitude in Sep/Oct and Apr/May, respectively. The δ^15^N of POM was substantially higher in Sep/Oct, with an average of 1.5 ± 1.0 ‰ (Fig. 5 c). In Apr/May, values were lower (mean: 0.2 ± 1.3 ‰) and increased from the north (station 3: -1.0 ‰) to the south (station 9: 2.7 ‰; Fig. 5 d).

The patterns in zooplankton abundance and δ^13^C were different from those observed in phytoplankton and POM (Fig. 5 e, f). The zooplankton abundances were up to one order of magnitude higher in Sep/Oct versus Apr/May, whereby this temporal change was less pronounced in the north and center of the lake. In Sep/Oct, zooplankton abundances reached a lake-wide maximum in the south (29.9 and 1.9 · 10^3^ ind. m^-2^ for the 95 µm and 250 µm size fractions, respectively) and a secondary peak of large zooplankton (250 µm)

in the northern part of the lake (1.3 ⋅ 10^3^ ind. m^-2^). In Apr/May, the abundances from both net types varied only slightly across the lake, reaching up to 2.0 and 0.5 · 10^3^ ind. m^-2^ for the 95 µm and 250 µm size fractions, respectively (Fig. 5 f). Zoo- and phytoplankton abundances were negatively correlated in Sep/Oct, i.e. phytoplankton (>10 µm) decreased, when zooplankton increased (*rho* = -0.58, *p* ∼ 0.1, Spearman’s rank correlation). No significant correlation between phyto- (>10 µm) and zooplankton was found in Apr/May (*rho* = 0.38, *p* > 0.3, Spearman’s rank correlation).

The zooplankton δ^13^C values changed slightly between the seasons. In Apr/May, both size fractions were ∼0.6 ‰ heavier compared to zooplankton from Sep/Oct, opposite of what we found in POM. Overall, zooplankton δ^13^C lie between -24.7 and -21.1 ‰ (Fig. 5 g, h). In Sep/Oct, we observed a trend towards higher values in the south (Δ +1 ‰ for both net types), whereas the highest values in Apr/May occurred towards the northern (−22.1 ‰) and southern (−21.1 ‰) extremities of the lake, with a minimum of -24.0 ‰ in the central region. The zooplankton δ^15^N values tended to be higher in the 250 µm compared to the 95 µm fraction. Overall, they were similar between the campaigns, but showed generally lower values in the north (2.5-6.3 ‰) compared to the center and the south (3.2-10.4 ‰; Fig. 5 i, j).

The zooplankton community composition was dominated by copepods (Fig. S3). In Sep/Oct, cyclopoids dominated the small size fraction (on average 66 and 78 % for 25 and 95 µm, respectively), whereas calanoids were prevalent in the 250 µm fraction (mean: 74 %). In Apr/May, calanoid copepods dominated the zooplankton community throughout all size fraction, but the relative abundance of cyclopoids was highest in the small size fractions here, too (up to 78 % in the north basin). The freshwater jellyfish *Limnocnida tanganyicae medusa* appeared in higher relative abundances in Sep/Oct, especially in the 250 µm size fraction, reaching up to 19 % at station 7. The contributions of shrimps to the total zooplankton community was low in our samples, reaching its highest values in the south during both seasons (max. 16 %). Fish and insect larvae were rare in all our samples (<1 %).

Overall, we find that the phytoplankton abundances (>10 µm) decreased towards the south, whereas chlorophyll-*a* and zooplankton reached the highest values in the south. Zooplankton δ^13^C and δ^15^N were variable, but also showed the highest values in the south. The southward increase was more expressed in POM δ^13^C and δ^15^N. In addition, POM δ^13^C and δ^15^N were generally heavier in Sep/Oct compared to Apr/May.

### 3.4 Isotopic and elemental composition of bivalve and fish populations

#### 3.4.1 Variations in δ^13^C and δ^15^N

The δ^13^C of *P. spekii* revealed a consistent seasonal and latitudinal dynamic (Fig. 6 a, b). We found a significant divergence between northern and southern populations in Sep/Oct (mean -22.5 and -21.4 ‰, respectively; Mann-Whitney U test, *p* < 0.01), whereas the δ^13^C values of the *P. spekii* samples from the two basins overlapped in Apr/May (means of -22.7 and -22.3 ‰, respectively), with northern samples nesting fully within the range of southern samples. The samples from the northern basin showed little variation between the seasons, whereas samples collected in the southern basin differed significantly (Mann-Whitney U test, *p* < 0.001).

**Fig. 6:**
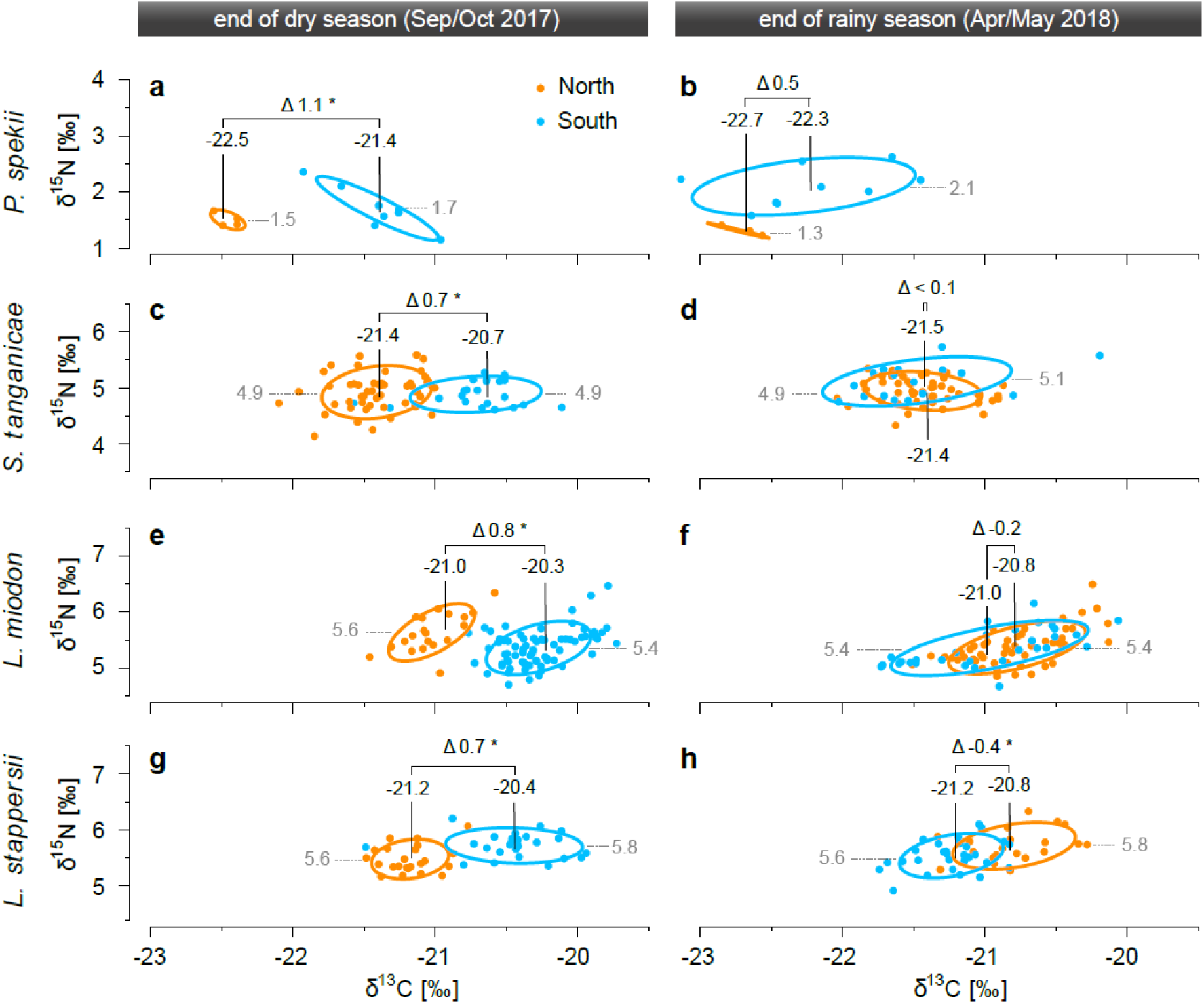
Carbon (normalized for C:N mass ratio according Post et al. (54)) and nitrogen stable isotope signatures of the major pelagic food web members. (**a,b**) the bivalve *Pleiodon spekii* as well as the fish (**c,d**) *Stolothrissa tanganicae*, (**e,f**) *Limnothrissa miodon*, and (**g,h**) *Lates stappersii* at the end of the dry season (left) and the end of the rainy season (right). Orange dots represent the northern basin (stations 1 and 2) and blue dots represent the southern basin (stations 7 and 9). Numbers indicate the mean δ^13^C (black) and δ^15^N (grey) of a population and Δ denotes the δ^13^C difference between the southern and northern populations. Significant differences in δ^13^C are marked by stars. Ellipses encompass approximately 95 % of the data of each population. Note different y-axis limits.

In Sep/Oct, the mean δ^13^C values of the populations of the largely planktivorous sardines *Stolothrissa* and *Limnothrissa* as well as the zooplankti- and piscivorous *Lates stappersii* diverged significantly (Mann-Whitney U tests, *p* < 0.001) by approximately 0.7 ‰ between the north and the south (Fig. 6 c, e, g). By contrast, the differences in mean values were completely erased or going slightly into the opposite direction in Apr/May (Fig. 6 d, f, h).

Moreover, the southern populations of all three species exhibited the highest δ^13^C with mean values of -20.7 ‰, -20.3 ‰, and -20.4 ‰ for *Stolothrissa*, *Limnothrissa,* and *Lates stappersii*, respectively. By contrast, the δ^13^C of the northern population in Sep/Oct was similar to the population-wide averages from both basins in Apr/May and never differed by more than 0.4 ‰. The lowest mean values were consistently observed in Apr/May (−21.5 ‰, -21.0 ‰, and -21.2 ‰ for *Stolothrissa*, *Limnothrissa,* and *Lates stappersii*, respectively).

In contrast to *P. spekii* and *Stolothrissa*, the southern populations of *Limnothrissa* and *Lates stappersii* showed 0.2 and 0.4 ‰ lower δ^13^C values compared to the northern one in Apr/May, respectively. This difference was not significant for *Limnothrissa* (Mann-Whitney U test, *p* > 0.05) and was significant for *Lates stappersii* (Mann-Whitney U test, *p* < 0.001). It is worth noting that samples of *Limnothrissa* and *Lates stappersii* from Apr/May included in our analysis were slightly unbalanced, with samples from the north being larger compared to samples from the south (Fig. S2). Since larger individuals in these two species tend to have less depleted δ^13^C values (Fig. S4 c, e), the δ^13^C values of the northern populations, and with them the difference between the basins, are slightly overestimated here.

We had fewer samples of the large predators *Lates microlepis*, *Lates mariae*, and *Lates angustifrons*, preventing an in-depth statistical analysis, but the results hint at similar patterns. Across our entire data set, these three species showed the highest δ^13^C values, with most observations being heavier than -21 ‰ (Figs. S4 g, i. k; S5). In Sep/Oct, the δ^13^C of both *Lates microlepis* and *Lates angustifrons* specimens were >0.5 ‰ lighter than the individuals from the southern populations. In Apr/May, samples from both the north and south basins were only available for *Lates mariae*. Here, the δ^13^C values from both basins varied within the same range and their averages differed only slightly (north: -20.1‰; south: -20.3 ‰).

Similar to zooplankton and POM *P. spekii* showed consistently higher δ^15^N values in the southern basin, reaching 1.7 and 2.1 ‰ on average in Sep/Oct and Apr/May, respectively, compared to the northern basin with averages of 1.5 and 1.3 ‰, for the two campaigns (Fig. 6 a, b). In contrast to δ^13^C, we observed no systematic seasonal or regional differences in fish δ^15^N (Figs. 6 c-h; S5). *Stolothrissa* exhibited the lowest values with population averages spanning from 4.9 to 5.1 ‰. The other sardine, *Limnothrissa*, had markedly higher values (means: 5.4-5.6 ‰). Most observations ranged between 4-6 ‰ and 4.5-6.5 ‰ for the two species, respectively (Fig. S4 b, d), whereby the variability appeared to be neither related to site nor season, and only to a small extent to size. By contrast, the δ^15^N of the larger *Lates* species were primarily related to size (Fig. S4 f, h, j, l). Individuals smaller than 200-250 mm showed an increase from <4 ‰ up to around 6-8 ‰, which then flattened out at these high values for specimens >250 mm. In the >250 mm size class, the δ^15^N of *Lates stappersii* and *Lates microlepis* never exceeded 7.5 ‰, whereas *Lates mariae* and *Lates angustifrons* reached values of >8 ‰ (Fig. S4 f, h, j, l). None of the *Lates* species revealed clear population-wide differences in δ^15^N between the basins or seasons (Figs. 6 g, h; S5).

Overall, we find that the different fish species and *P. spekii* revealed congruent patterns in δ^13^C with diverging northern and southern populations when upwelling/mixing in the south were strongest, i.e. at the end of the dry season in Sep/Oct. By contrast, δ^13^C values converged, and were generally lower, when the lake was more heavily and homogenously stratified, i.e. at the rainy season-dry season transition (Apr/May). On the contrary, the fish populations exhibited no clear basin-scale trends in δ^15^N, whereas *P. spekii* showed slightly higher values in the south during both seasons.

#### 3.4.2 C:N ratios and estimated lipid content

We observed strong changes in C:N ratios between Sep/Oct and Apr/May at lower and middle trophic levels, but not at high trophic levels; there were no clear differences between north and south in all organisms (Fig. 7). *Pleiodon spekii* and zooplankton showed consistently lower C:N ratios in Sep/Oct compared to Apr/May (Fig. 7 a, c, e), but the low sample size (n ≥ 3) ruled out a statistical analysis. We observed the largest seasonal change in zooplankton (250 µm) from the southern basin. The medians differed between 5.1 (Sep/Oct) and 6.8 (Apr/May). The differences were smaller in *P. spekii*, revealing an overall minimum median of 3.9 in the northern basin in Sep/Oct and a maximum of 4.3 in the southern basin in Apr/May.

**Fig. 7:**
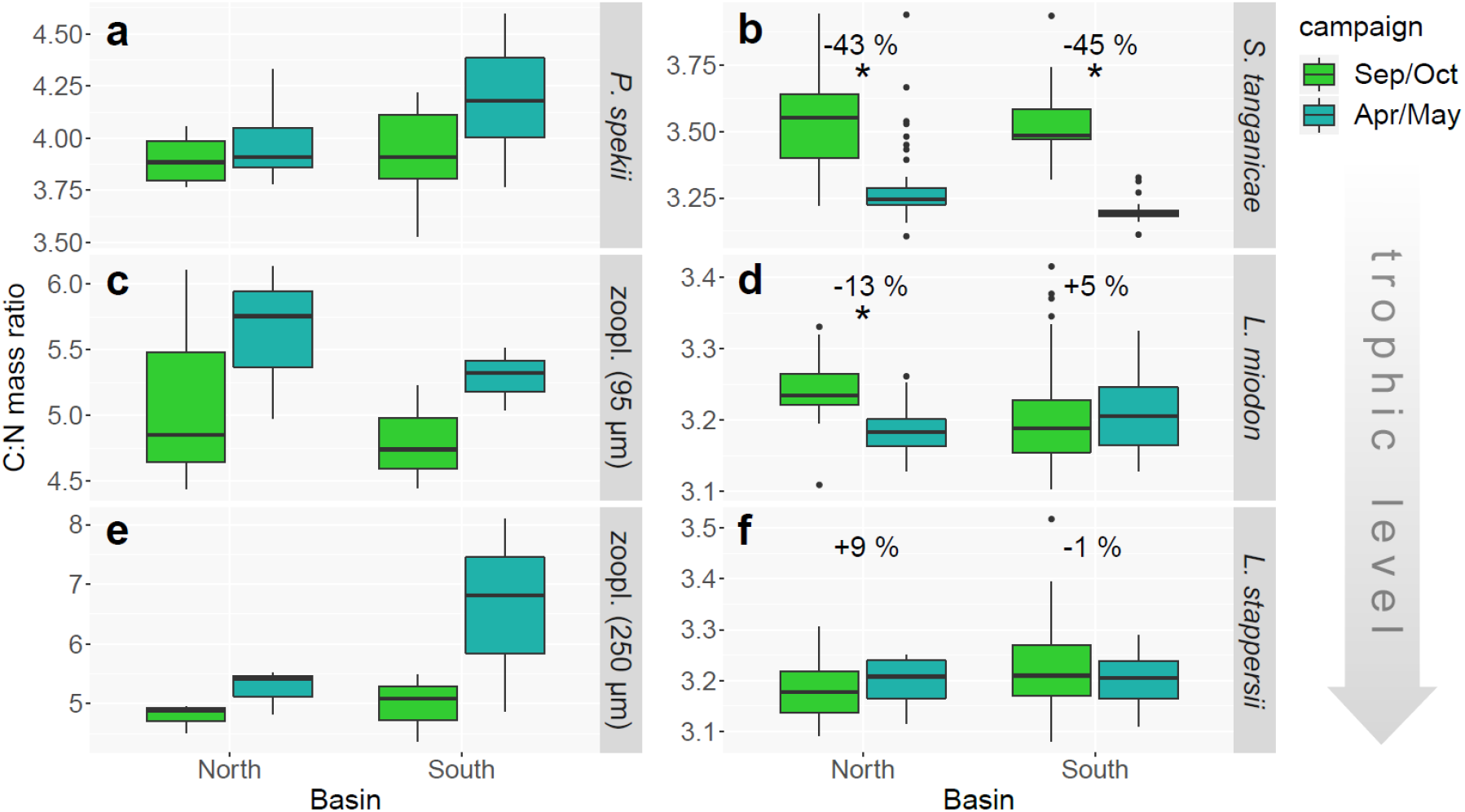
Mass C:N ratios of primary consumers (left) and fish tissue (right) for the different sampling campaigns and basins of Lake Tanganyika. (**a**) *Pleiodon spekii*, (**c**, **e**) zooplankton, (**b**) *Stolothrissa tanganicae*, (**d**) *Limnothrissa miodon*, and (**f**) *Lates stappersii*. Stars indicate significant differences between campaigns and numbers depict the % change in estimated lipid content according to Post et al. (54). Note varying y-axis scaling.

Contrary to the trends in *P. spekii* and zooplankton, the sampled fish species showed a tendency towards higher C:N ratios in Sep/Oct (Fig. 7 b, d, f). This trend vanished with increasing trophic level of the fish species. The C:N ratios of the planktivorous clupeid *Stolothrissa* decreased significantly (Mann-Whitney U tests, *p* < 0.001) between Sep/Oct and Apr/May in both the northern and southern basins (medians of 3.65 versus 3.24 and 3.49 versus 3.19, respectively). *Limnothrissa* revealed significant changes in the north (Sep/Oct median: 3.23; Apr/May median: 3.18; Mann-Whitney U test, *p* < 0.001), whereas the pattern was reversed and insignificant in the south (Sep/Oct median: 3.19; Apr/May median: 3.21; Mann-Whitney U test, *p* > 0.1). By contrast, the C:N ratios of *Lates stappersii* varied only within a narrow range, with medians spanning from 3.18 to 3.21 and no significant differences between seasons (Mann-Whitney U tests, *p* > 0.1).

Using the model from Post et al. (54), we estimated the lipid contents in the dorsal muscle tissue of the investigated fish species from their C:N ratios. This analysis suggests a reduction of lipid content in *Stolothrissa* from Sep/Oct to Apr/May by 43 and 45 % in the north and south, respectively. *Limnothrissa* exhibited a 13 % decrease (north) and a 5 % increase (south). *Lates stappersii* showed the lowest differences with a 9 % increase (north) and 1 % decrease (south). A gravimetric determination of lipid content from selected *Stolothrissa* samples confirmed that higher C:N ratios translate into higher lipid contents (linear regression, *R^2^* = 0.91, *p* < 0.01, *n* = 5; Fig. S6).

## 4 Discussion

### 4.1 Effect of upwelling and mixing on the isotopic composition of the planktonic food web

Upwelling and convective mixing moderate the transport of nutrients to the surface waters, and thus drive biological productivity during the dry season in Lake Tanganyika. In this study, we compare two contrasting hydrodynamic situations: First, the period of re-establishing water column stratification at the end of the dry season (Sep/Oct). During this time, stratification was weaker and the thermocline was still absent at the southernmost station, enabling particularly high nutrient fluxes in the south. Second, the period of lake-wide stratification at the rainy season-dry season transition (Apr/May).

Here, the water column experienced stronger stratification from the preceding rainy season and beginning trade winds initiated the upwelling in the south, resulting in overall lower nutrient fluxes with a maximum in the south. Our data show that upwelling and mixing do not only influence biological productivity, but also impact the isotopic composition of all food web members, likely owing to differences in primary productivity and N acquisition pathways of phytoplankton. Such systematic changes can be used to delineate regional fish populations.

In contrast to the southward increase in nutrient availability, we detected a southward decrease in the abundance of large-sized phytoplankton (>10 µm) a pattern that has previously been observed in Lake Tanganyika (70), pointing out additional ecological controls, such as zooplankton grazing or competition within the phytoplankton community. In Lake Tanganyika, the nano- and pico size fractions (<10 µm) are more competitive under nutrient-rich conditions and therefore dominate the phytoplankton community in south (71, 72). As a result of their high densities, total phytoplankton abundance and biomass is generally highest during the dry season upwelling in the south (18,21,73).

In addition, upwelling/mixing and the subsequent stimulation of primary productivity, place important bottom-up control on the abundances of zooplankton in Lake Tanganyika. This was evidenced by high zooplankton abundances in the dry season with maxima in the southern basin (Fig. 5; 18,70), which in turn sustain the growth of the pelagic fish populations, especially the sardine species (4,23,24,75). The high zooplankton abundances can in turn exert top-down control over phytoplankton, which is indicated by the negative correlation between the phytoplankton (>10 µm) and zooplankton abundances in Sep/Oct. This grazing effect may have also been responsible for the absence of strong differences in chlorophyll-*a* between Sep/Oct and Apr/May. Zooplankton abundance, and thus grazing pressure, exhibited no clear latitudinal trend during the April/May campaign, when the lake-wide stratification was stronger, and the north-south gradients in nutrient availability and biological productivity were not as pronounced.

The varying hydrodynamics were also associated with distinct δ^13^C-POM signatures that may reflect differences in primary productivity. The average δ^13^C was ∼5 ‰ heavier in Sep/Oct compared to Apr/May, and the δ^13^C increased by ∼2 ‰ from north to south during both campaigns (Fig. 5 c, d). In Lake Tanganyika, previous studies also revealed heavier δ^13^C-POM values in the dry season (51–53), even though the differences were smaller (max. 3.1 ‰) than in our study (max. 6.7 ‰ ), possibly due to the varying timing of the sampling. These differences in δ^13^C-POM likely reflect the well-documented changes in primary production (19,73,76), where heavier isotopic signatures in POM mirror the incorporation of a larger ^13^C fraction by higher photosynthesis and cell growth rates (44, 45) and a stronger drawdown of the DIC pool (46–48,77). The links between stratification, vertical nutrient supply, and primary productivity in Lake Tanganyika are well established (10–12,19) and several studies have used the δ^13^C of sediment POM to infer primary productivity (7, 12). Our own CO_2_ fixation rate measurements, done during Apr/May, show evidence for higher productivity rates in the southern basin at station 7 in the south compared to station 2 in the north (Fig. 4). On the other hand, upwelling of intermediate waters will not only supply nutrients, but also isotopically light DIC (depleted by ∼1 ‰; Fig. 3; 59,75). Although this mechanism will slightly dilute ^13^C enrichment, our proposed mechanism of higher primary productivity is apparently strong enough to overcome this depletion in δ^13^C-POM, ultimately leading to higher δ^13^C-POM when upwelling/mixing is stronger.

The analogous pattern in δ^15^N-POM implies that upwelling and mixing may have influenced the N sources of primary producers, where lighter values are typically interpreted as inputs from N fixation (77, 79). POM δ^15^N values increased from -1.0 ‰ at station 3 to 2.7 ‰ at station 9 in the south in Apr/May, concurrent with a decrease in filamentous, N-fixing cyanobacteria (56), whereas it fluctuated with slightly higher values (−0.3-2.6 ‰) in Sep/Oct devoid of a latitudinal or phytoplankton composition related pattern. We also observed no correlation between the presence of surface nitrate and δ^15^N-POM, which may have induced fractionation effects during nitrate-uptake.

When free nitrate remains, the phytoplankton community does not represent a complete sink of the upward diffusing nitrate, i.e. the residual nitrate should be isotopically heavy and phytoplankton relatively light. In line with the higher density of N-fixing cyanobacteria (21,56,72), the generally lighter δ^15^N-POM in Apr/May (Δ-1.4 ‰) point at higher inputs from N fixation compared to Sep/Oct, when nutrient fluxes are higher due to upwelling/mixing.

The isotopic composition of the zooplankton community did not show clear latitudinal and seasonal patterns, pointing at additional influencing factors than the signal from the base of the food web, i.e. POM. The δ^13^C values in our zooplankton samples oscillated between -24.7 and -21.1 ‰, in accordance with previous isotope surveys. The δ^15^N values from the north were also in agreement with other studies, whereas the maxima in the south, where previously no isotopic characterization of the food web was undertaken, exceeded earlier reports by min. 2.7 ‰ (Fig. 5; 45–47,62). The high intra-basin variability in δ^13^C and δ^15^N as well as the high absolute values relative to other members of the food web, with some zooplankton δ^15^N exceeding top predator fish δ^15^N values, may be in part attributable to varying zooplankton community compositions (52, 68).

Such values from a pooled zooplankton sample are not unexpected, because zooplankton communities consist usually of members from several trophic levels (e.g. 46,77,78), and our samples represent batch samples from entire zooplankton communities formed by many different species, genera, and families. In addition, the zooplankton community is notoriously hard to sample and standard netting techniques do not capture fast swimmers such as shrimps efficiently, therefore often underestimating their abundances (26).

Indeed, shrimps only made up minor proportions in our samples (Fig. S3) and previous work showed that they have δ^13^C and δ^15^N values lower than our community isotope values (52,53,68). However, reported δ^15^N values of individual zooplankton taxa, including detrivorous jellyfish and fish larvae, do not exceed 5.9 ‰ (52,53,68) and therefore fail at explaining the high δ^15^N in our measured community isotope samples from the south (>10 ‰). In line with our results, earlier reports of bulk community samples found high δ^15^N values between 6 and 8 ‰ (51), raising questions about the utility of using bulk community samples. Combining the taxonomic assessment of the community with an isotopic characterization of individual zooplankton taxa would thus be valuable in future food web studies.

In summary, our results point to a pivotal role of nutrient upwelling and mixing for sustaining the high biological productivity in the south basin during the dry season.

Upwelling-related increases in primary productivity and decreases in N fixation likely resulted in markedly heavier planktonic δ^13^C and slightly heavier δ^15^N values in the south (Fig. 8 a). The slightly higher zooplankton δ^13^C and δ^15^N values in the southern basin may reflect the isotopic imprint of the upwelling/mixing, but a clear north-south trend may be masked to some extent by concomitantly shifting community composition effects.

**Fig. 8:**
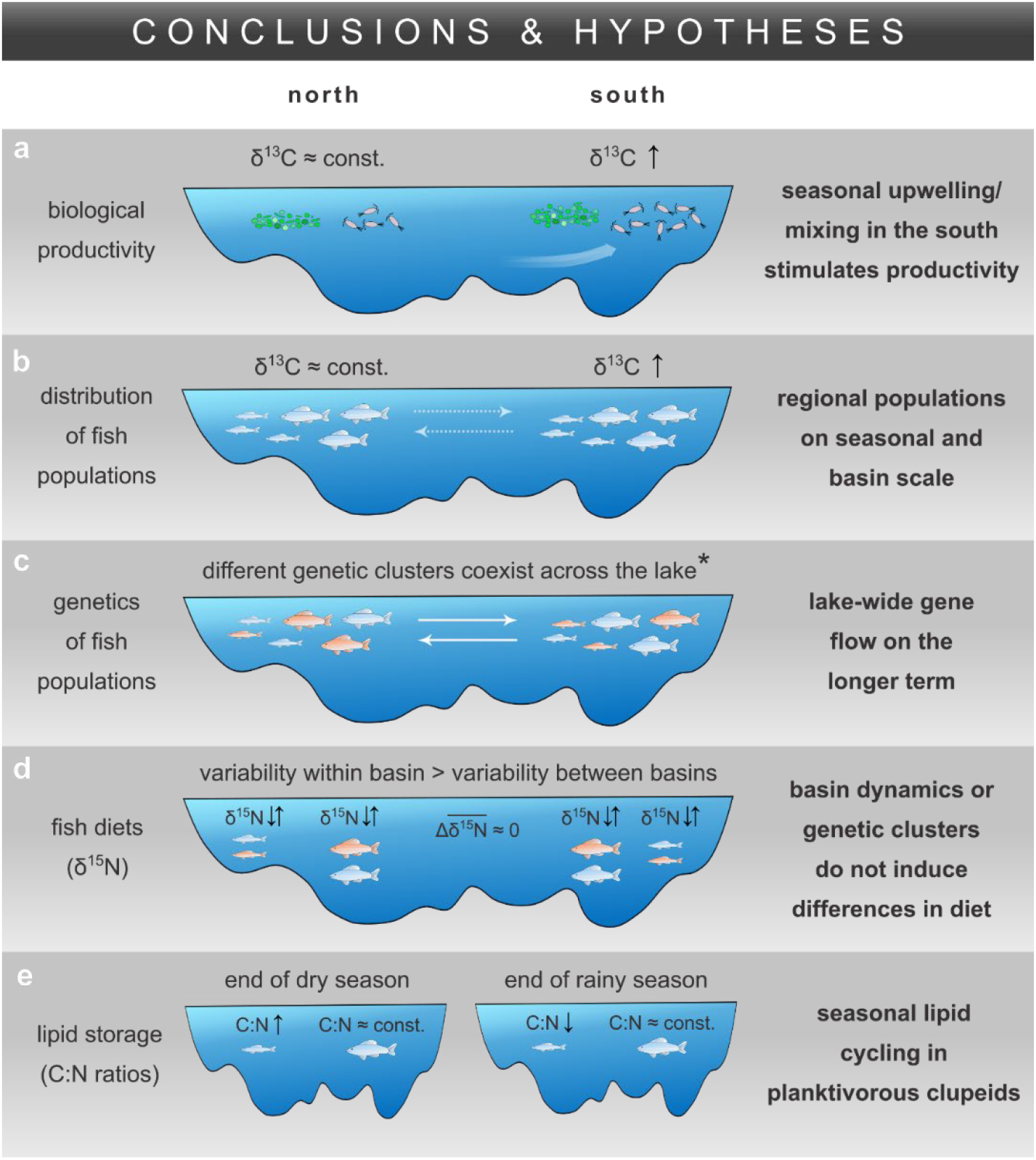
Sketch synthesizing the main conclusions and hypotheses of the study. (**a**) Biological productivity of phyto- and zooplankton based on abundance and δ^13^C data, which were used to (**b**) infer the distribution of regional fish populations from their δ^13^C signatures. (**c**) The regional isolation of the fish populations is apparently insufficient to suppress lake-wide gene flow. (**d**) The regional fish populations as well as different genetic clusters did not exhibit systematic differences in δ^15^N. (**e**) The clupeid *Stolothrissa* exhibited strong seasonal changes in C:N, i.e. lipid content, indicating lipid storage after the productive dry season. * results from Junker et al. (35) and Rick et al. (36)

### 4.2 Isotopic imprints from upwelling and mixing reveal regional fish populations

The seasonally and regionally varying hydrodynamic conditions also determine the C isotopic compositions of organisms higher in the food web, through the incorporation of phyto- and zooplankton prey. Due to the longevity of many organisms higher in the food web, they integrate the isotopic signals from their food over a longer time span. For instance, tissue turnover in bivalves is significantly slower than in phyto- and zooplankton (51, 82). Therefore, we used the filter-feeding bivalve *P. spekii* as a reference organism for the seasonal phytoplankton isotopic signals. *Pleiodon spekii* lives for at least five years at a stationary location and was successfully used to record upwelling events in Lake Tanganyika (83). We find that the southern *P. spekii* samples in Sep/Oct diverge significantly in δ^13^C from northern samples (Δ1.1 ‰). Lake-wide stratification over the rainy season on the other hand resulted in converging δ^13^C values between samples from both basins in Apr/May (Fig. 6 a, b).

Such seasonal cycles in δ^13^C signals are common in aquatic food webs and propagate along the trophic chain (50, 84). The typical muscle half-life ranging from a few weeks in juvenile fish to several months in adults explains the incorporation of seasonal dietary isotope patterns (85–87). Accordingly, we find similar patterns as in *P. spekii* in the three major pelagic fish species of Lake Tanganyika, with a latitudinal difference of ∼0.7 ‰ heavier fish tissue δ^13^C in the south (Fig. 6). The aligning δ^13^C values between northern and southern samples during the rainy season are again in agreement with the more uniform primary productivity patterns. Our results point to fish populations confined to regional foraging grounds in the respective basins, which therefore record the latitudinal isotope gradients (Fig. 8 b).

However, our previous high resolution population genetic work did not find evidence for genetic differentiation between the north and south basins in any of the six fish taxa investigated in this study. Instead, populations in the north and south basins are closely related (34–36). The limited genetic differentiation in these species is not spatially restricted, with the exception of a case in *Lates mariae*. In Rick et al. (36), we found one *Lates mariae* cluster confined to the extreme south end of the lake with strong genetic differentiation from individuals elsewhere in the south basin or in the rest of the lake.

Thus, the genetic structure of the fish populations cannot be explained by the basin-scale dynamics. This implies that the degree of geographical isolation between north and south basins itself is insufficient to suppress lake-wide gene flow in these pelagic fish species. In other words, the isotopically distinct fish populations can only be regarded as regional on rather short seasonal to multiannual time scales (Fig. 8 c).

### 4.3 Regional fish populations show no differences in δ^15^N or C:N ratios

Despite the absence of pronounced spatial genetic structure in either of the sardine (34, 35) or *Lates* species (36), regional fish populations may exhibit phenotypic changes in diet or lipid content in response to regionally different environments, which include a northern region with a more stable and clear water column and a plankton-rich upwelling region in the south (15,18,75).

The rather constant average δ^15^N of the studied fish (deviation < 0.2 ‰) and *P. spekii* (deviation < 0.8 ‰) among the sampling campaigns and basins indicates no strong differences in trophic level between the populations in the different lake basins. However, the difficulty of quantifying the trophic position of the fish species was exacerbated by the small differences between the trophic levels. Moreover, the zooplankton community δ^15^N exhibited strong intra-basin variability, with maxima similar to the highest fish δ^15^N values, which raises doubt about the usefulness of comparing bulk community with tissue samples (88). Both the high variability in δ^15^N between individual zooplankton taxa (52,53,68) and unknown trophic discrimination factors (89), which appear to deviate from the norm in Lake Tanganyika (52, 68), further aggravate assessing subtle differences in the specific diets. Compound specific isotope analyses of amino acids may further help constraining the trophic relationships (90).

Nonetheless, on a basin-scale, the relatively consistent fish and *P. spekii* δ^15^N demonstrate that the isotopic composition of their N sources does not vary substantially throughout the year and among basins, whereas POM and zooplankton showed some tendency towards higher values in the south. Instead, larger δ^15^N variations of up to >2‰ among individuals of a similar size found at the same location and time indicate that other factors than the basin-scale hydrodynamics influence the diets of the studied fish taxa. We found no clear evidence, however, that differences in δ^15^N were linked to the different genetic clusters in *Limnothrissa* or the four *Lates* species (Figs. S7; 8 d).

Fish use lipids to store energy during times of abundant food supply to bridge resource limited periods (38, 39). However, in congruence with the absence of basin-scale genetic structure, we found no regional differences in C:N ratios as proxy for lipid content (Fig. S6; 48,49,88), but we did find seasonal changes: the smallest species, *Stolothrissa,* showed a significantly higher C:N ratio, i.e. lipid content, at the end of the productive dry season (Fig. 7), which translates to >40 % change in lipid content according to the model of Post et al. (54). Seasonal lipid cycling is expected to be more pronounced in smaller fish, due to higher metabolic rates (92) and their planktivorous diet. While predators, i.e. *Lates stappersii* and large *Limnothrissa*, feed on both fish and zooplankton, the solely planktivorous *Stolothrissa* must cope with the strong seasonal fluctuations in plankton productivity. Thus, *Stolothrissa* may have a life history adapted to building reserves during the productive dry season for the following rainy season, when resources are less abundant. Alternatively, the changing C:N may relate to spawning activities (93).

However, spawning peaks were reported to occur in September and April-July (94, 95), i.e. during both our sampling occasions (Sep/Oct and Apr/May), and can thus not explain the observed changes in C:N between those two time points. The seasonal effect was less pronounced in the slightly larger *Limnothrissa* and was clearly absent in *Lates stappersii*, possibly due to their larger sizes and more piscivorous diets (Fig. 8 e). Overall, the δ^15^N and C:N values indicate similar diets and lipid contents of the northern and southern fish populations. We hypothesize that the long term gene flow across the lake may inhibit the development of ecological differences among the regional fish populations in response to the basin-scale environmental conditions.

## 5 Conclusions

In this study, we showed that the seasonal upwelling and mixing in the south basin of Lake Tanganyika induce distinct isotopic imprints at the primary producer level. These distinct isotopic signals can be tracked across the entire pelagic food web. Using δ^13^C as tracer, we identified fish populations with regional foraging grounds, implying some degree of isolation on a seasonal and basin-wide scale. Correspondingly, regional fishery management strategies may include basin-scale quotas. Our elemental and bulk isotopic composition data provide no clear evidence for strong physiological or dietary differences among these regional populations. Although lake-wide gene flow may inhibit the evolution of clear ecological differences between the regional populations, a different set of methods may reveal some ecological variation that cannot be resolved with bulk elemental and isotopic analyses. In the context of assessing the vulnerability of Lake Tanganyika’s pelagic food web in a warming climate, our study indicates that the economically relevant pelagic fish species are genetically adapted to the whole lake although they form regional populations at the seasonal time scale.

## Acknowledgements

We are grateful for the support from our research collaborators at the Tanzania Fisheries Research Institute, particularly the Directors Rashid Tamatamah and Semvua Mzighani as well as Mary Kishe. Special thanks go to Mupape Mukuli as well as the captain and crew of the *M/V Maman Benita* for their steady toil in organizing and conducting the cruise work with us. We also thank Andreas Brand, Kathrin B.L. Baumann, and Tumaini M. Kamulali for their help during field work, Serge Robert and Fabian Kuhn for assistance in the lab, and Eliane Scharmin for administrative support. Special thanks go to Jessica A. Rick for providing help in the field, the data of the *Late*s genetic clusters, and comments on the manuscript. Thanks to Blake Matthews for insightful discussions. This work was funded by the Swiss National Science Foundation (grant CR23I2-166589). Thanks to the Tanzania Commission for Science and Technology (COSTECH) for granting the research permits.

## Author contributions

**Conceptualization:** Benedikt Ehrenfels, Julian Junker, Ismael A. Kimirei, Ole Seehausen, Catherine E. Wagner, Bernhard Wehrli

**Data Curation:** Benedikt Ehrenfels, Julian Junker, Demmy Namutebi, Cameron M. Callbeck, Anthony Kalangali, Athanasio S. Mbonde

**Formal Analysis:** Benedikt Ehrenfels, Julian Junker, Demmy Namutebi, Cameron M. Callbeck

**Funding Acquisition:** Julian Junker, Ismael A. Kimirei, Carsten J. Schubert, Ole Seehausen, Catherine E. Wagner, Bernhard Wehrli

**Investigation:** Benedikt Ehrenfels, Julian Junker, Cameron M. Callbeck, Christian Dinkel, Anthony Kalangali, Ismael A. Kimirei, Athanasio S. Mbonde, Julieth B. Mosille, Emmanuel A. Sweke

**Methodology:** Benedikt Ehrenfels, Julian Junker, Christian Dinkel, Anthony Kalangali, Ismael A. Kimirei, Athanasio S. Mbonde, Julieth B. Mosille, Emmanuel A. Sweke, Carsten J. Schubert, Ole Seehausen, Catherine E. Wagner, Bernhard Wehrli

**Project Administration:** Benedikt Ehrenfels, Julian Junker, Christian Dinkel, Anthony Kalangali, Ismael A. Kimirei, Athanasio S. Mbonde, Julieth B. Mosille, Emmanuel A. Sweke, Ole Seehausen, Catherine E. Wagner, Bernhard Wehrli

**Resources:** Ismael A. Kimirei, Emmanuel A. Sweke, Carsten J. Schubert, Ole Seehausen, Catherine E. Wagner, Bernhard Wehrli

**Supervision:** Carsten J. Schubert, Ole Seehausen, Catherine E. Wagner, Bernhard Wehrli

**Validation:** Benedikt Ehrenfels, Julian Junker, Demmy Namutebi, Cameron M. Callbeck, Anthony Kalangali, Athanasio S. Mbonde, Julieth B. Mosille, Emmanuel A. Sweke

**Visualization:** Benedikt Ehrenfels

**Writing – Original Draft Preparation:** Benedikt Ehrenfels

**Writing – Review & Editing:** Benedikt Ehrenfels, Julian Junker, Cameron M. Callbeck, Ismael A. Kimirei, Carsten J. Schubert, Ole Seehausen, Catherine E. Wagner, Bernhard Wehrli

## Supporting information

**Fig. S1:** C:N mass ratio of (**a,b**) *Stolothrissa tanganicae,* (**c,d**) *Limnothrissa miodon*, and (**e,f**) *Lates stappersii* versus standard length in the northern and southern basins during the end of the dry season and the end of the rainy season. Only stations 1, 2 (north) and 7, 9 (south) are depicted. The shaded areas mark the 50 mm cut-off range for the population comparisons used in Fig. 5.

**Fig. S2:** C:N corrected (Post et al. 2007) δ^13^C of (**a,b**) *Stolothrissa tanganicae,* (**c,d**) *Limnothrissa miodon*, and (**e,f**) *Lates stappersii* versus standard length for the end of the dry season and the end of the rainy season. Only stations 1, 2 (north) and 7, 9 (south) are depicted. The shaded areas mark the 50 mm cut-off range for the population comparisons used in Fig. 4. The sampled populations of *L. miodon* and *L. stappersii* from the end of the rainy season (d,f) were characterized by dense clusters of observations within a narrow size and δ^13^C range which may have skewed the basin-scale comparisons.

**Fig. S3:** Zooplankton community compositions per station for various net types (25 µm, 95 µm, 250 µm) during end of the dry season (top) and the end of the rainy season (bottom).

**Fig. S4:** C:N corrected according to Post et al. (54) δ^13^C (left) and δ^15^N (right) of (**a**,**b**) *Stolothrissa tanganicae*, (**c**,**d**) *Limnothrissa miodon*, (**e**,**f**) *Lates stappersii*, (**g**,**h**) *Lates microlepis*, (**i**,**j**) *Lates mariae* and (**k,l**) *Lates angustifrons* versus standard length including all sampling locations and campaigns. Samples from the central basin and July 2017 were included for completeness, but were not included in the north-south and seasonal analysis presented in Fig. 6. Note the different y-axis scaling.

**Fig. S5:** Carbon (normalized for C:N mass ratio according to Post et al. (54)) and nitrogen stable isotope signatures of the large *Lates* species, namely (**a,b**) *Lates microlepis* (**c,d**) *Lates mariae*, and (**e,f**) *Lates angustifrons* at the end of the dry season (left) and the end of the rainy season (right). Orange dots represent the northern basin (stations 1-3) and blue dots represent the southern basin (stations 7-9). Numbers indicate the mean δ^13^C of a population. Only individuals >150 mm were included in this analysis to reduce ontogenetic effects on the isotope signatures.

**Fig. S6:** Lipid content versus C:N ratios of *Stolothrissa tanganicae*. Each dot represents a

## Notes

### Competing Interest Statement

The authors have declared no competing interest.

