## Supplementary information for "Isotopic signatures induced by upwelling reveal regional fish populations in Lake Tanganyika"

¶ shared first authorship


not peer-reviewed preprint

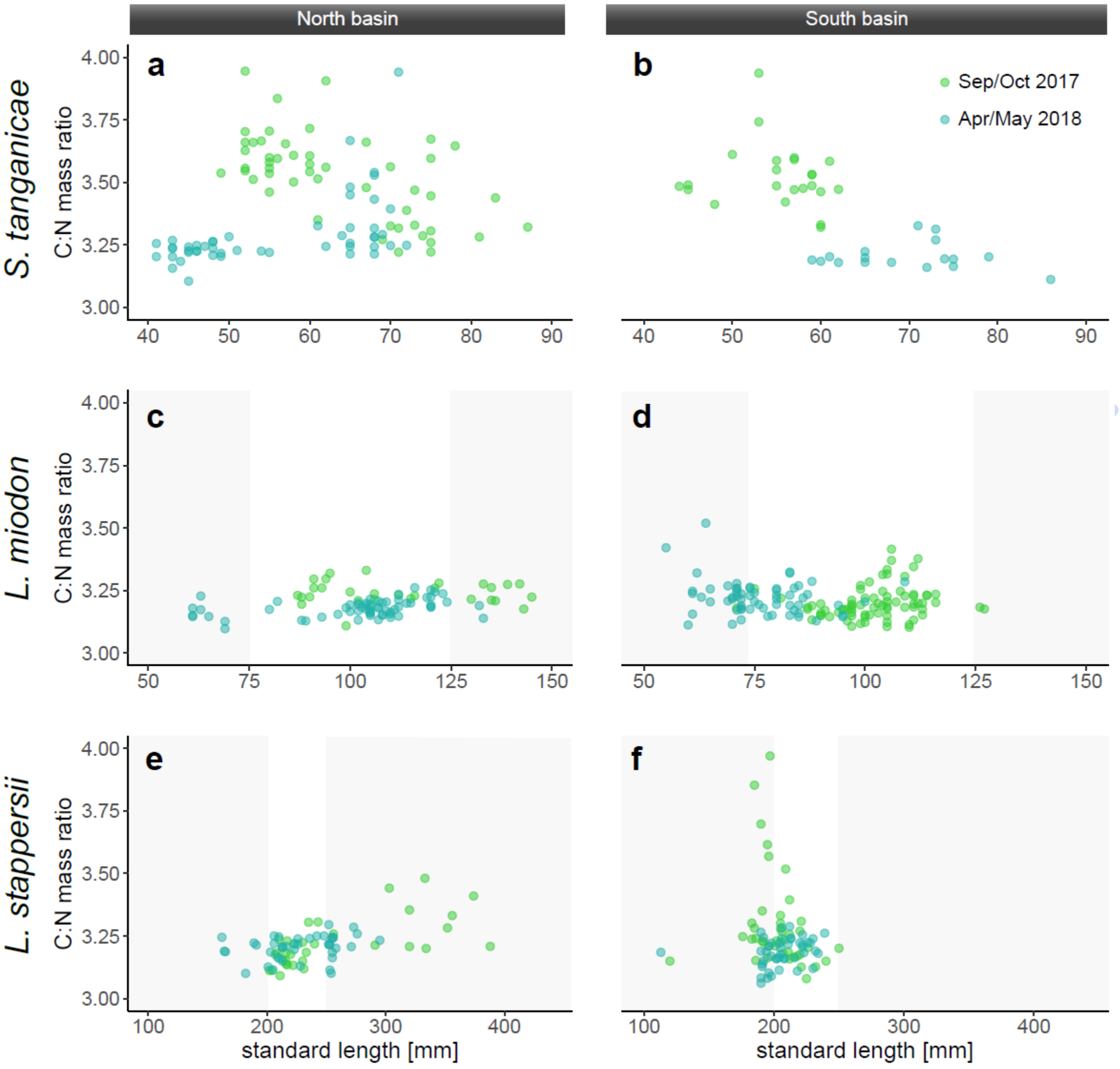


**Fig. S1:** C:N mass ratio of (**a,b**) *Stolothrissa tanganicae,* (**c,d**) *Limnothrissa miodon*, and (**e,f**) *Lates stappersii* versus standard length in the northern and southern basins during the end of the dry season and the end of the rainy season. Only stations 1, 2 (north) and 7, 9 (south) are depicted. The shaded areas mark the 50 mm cut-off range for the population comparisons used in Fig. 5.

not peer-reviewed

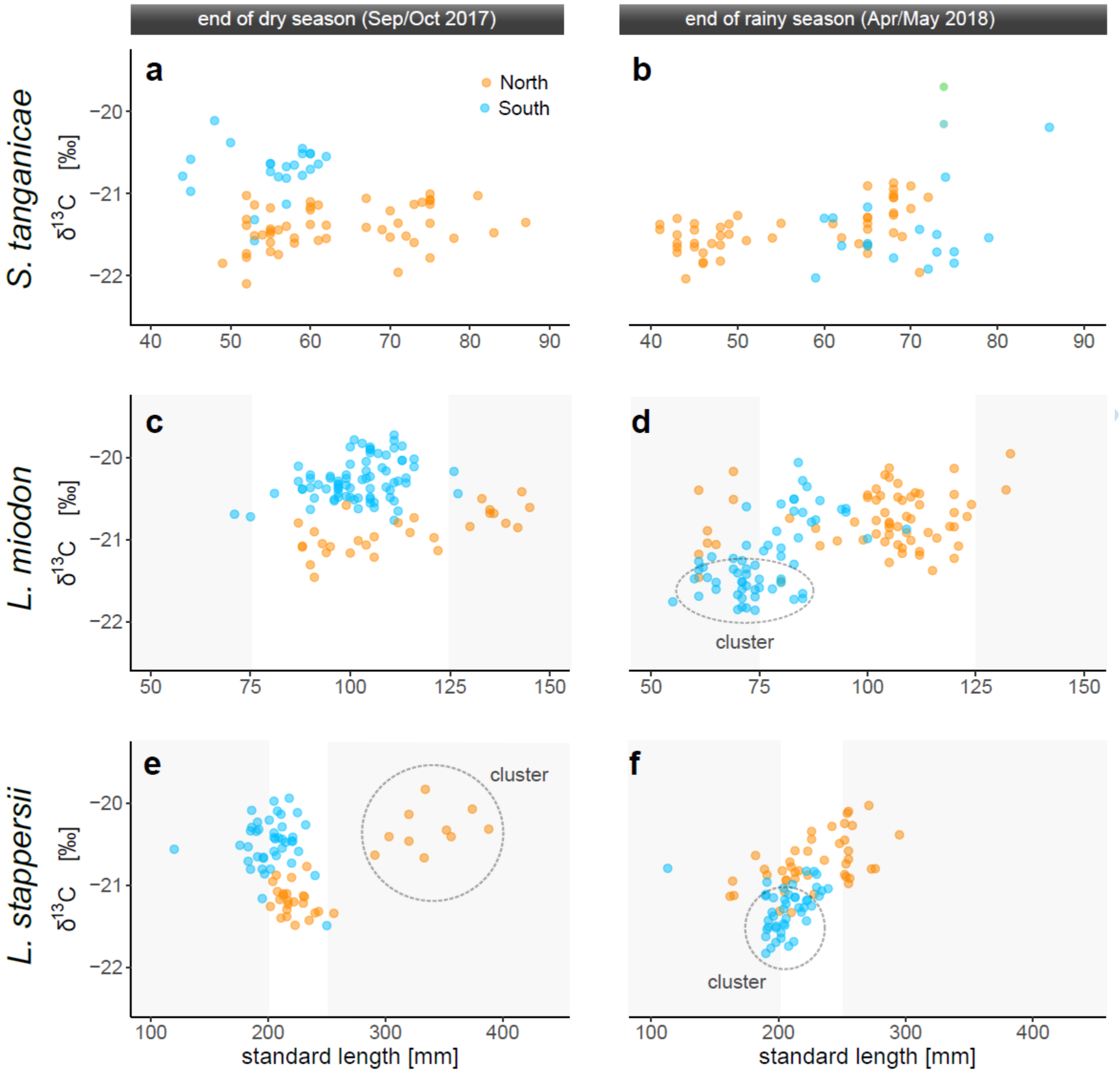


**Fig. S2:** C:N corrected (Post et al. 2007) $\delta^{13}C$ of (**a,b**) *Stolothrissa tanganicae,* (**c,d**) *Limnothrissa miodon*, and (**e,f**) *Lates stappersii* versus standard length for the end of the dry season and the end of the rainy season. Only stations 1, 2 (north) and 7, 9 (south) are depicted. The shaded areas mark the 50 mm cut-off range for the population comparisons used in Fig. 4. The sampled populations of *L. miodon* and *L. stappersii* from the end of the rainy season (d,f) were characterized by dense clusters of observations within a narrow size and $\delta^{13}C$ range which may have skewed the basin-scale comparisons.

not peer-

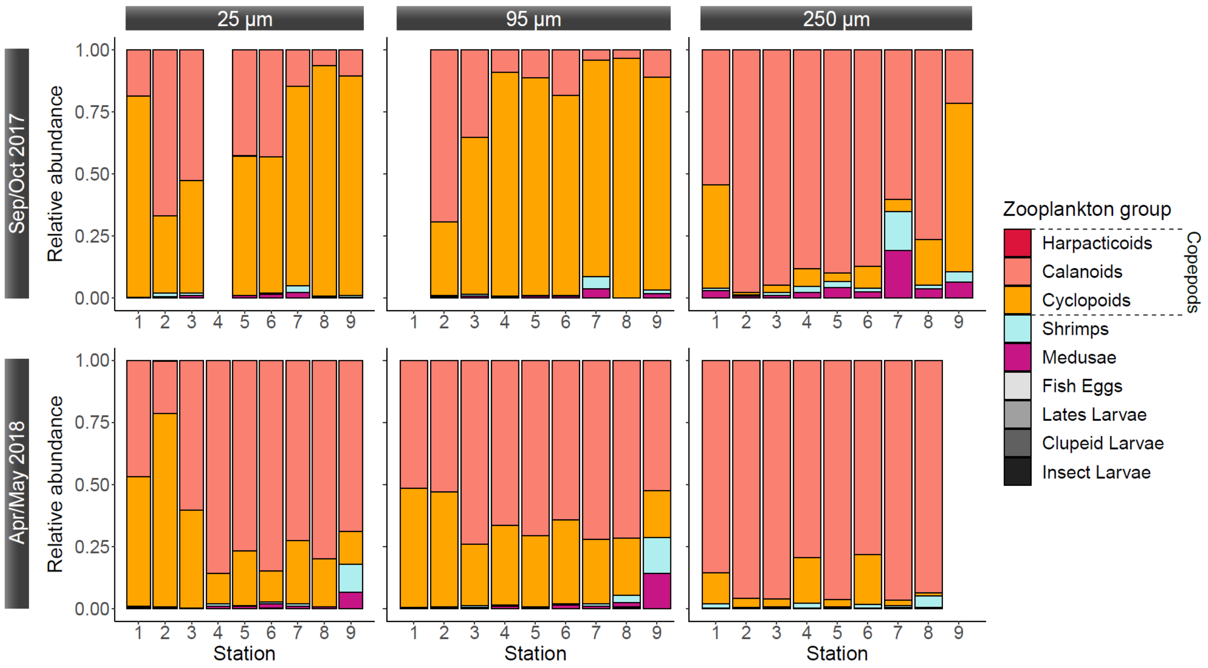


**Fig. S3:** Zooplankton community compositions per station for various net types (25 µm, 95 µm, 250 µm) during end of the dry season (top) and the end of the rainy season (bottom).

not peer-reviewed pr

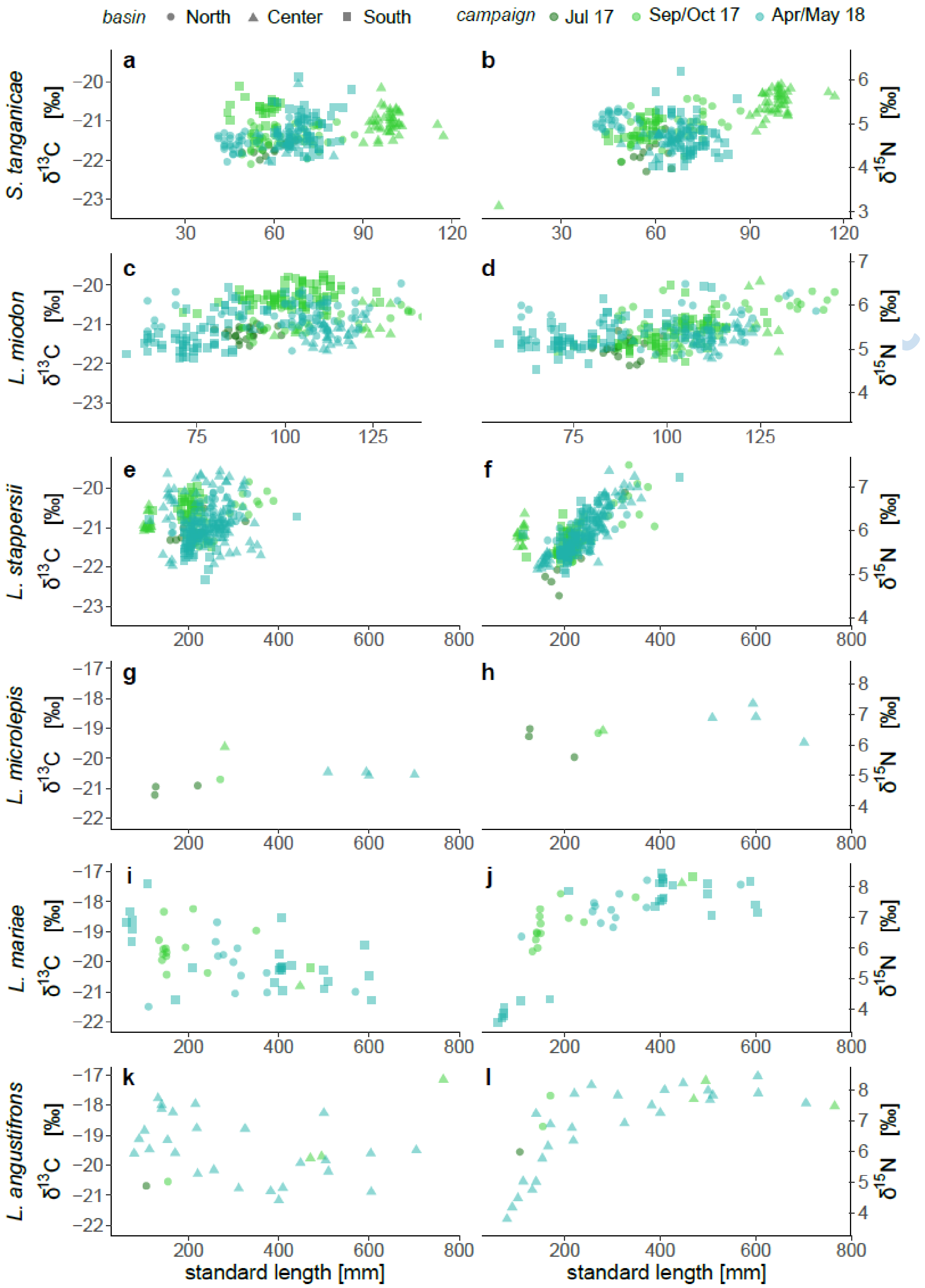


**Fig. S4:** C:N corrected (Post et al. 2007) $\delta^{13}C$ (left) and $\delta^{15}N$ (right) of (**a**,**b**) *Stolothrissa tanganicae*, (**c**,**d**) *Limnothrissa miodon*, (**e**,**f**) *Lates stappersii*, (**g**,**h**) *Lates microlepis*, (**i**,**j**) *Lates mariae* and (**k**,**l**) *Lates angustifrons* versus standard length including all sampling locations and campaigns. Samples from the central basin and July 2017 were included for completeness, but were not included in the north-south and seasonal analysis presented in Fig. 6. Note the different y-axis scaling.

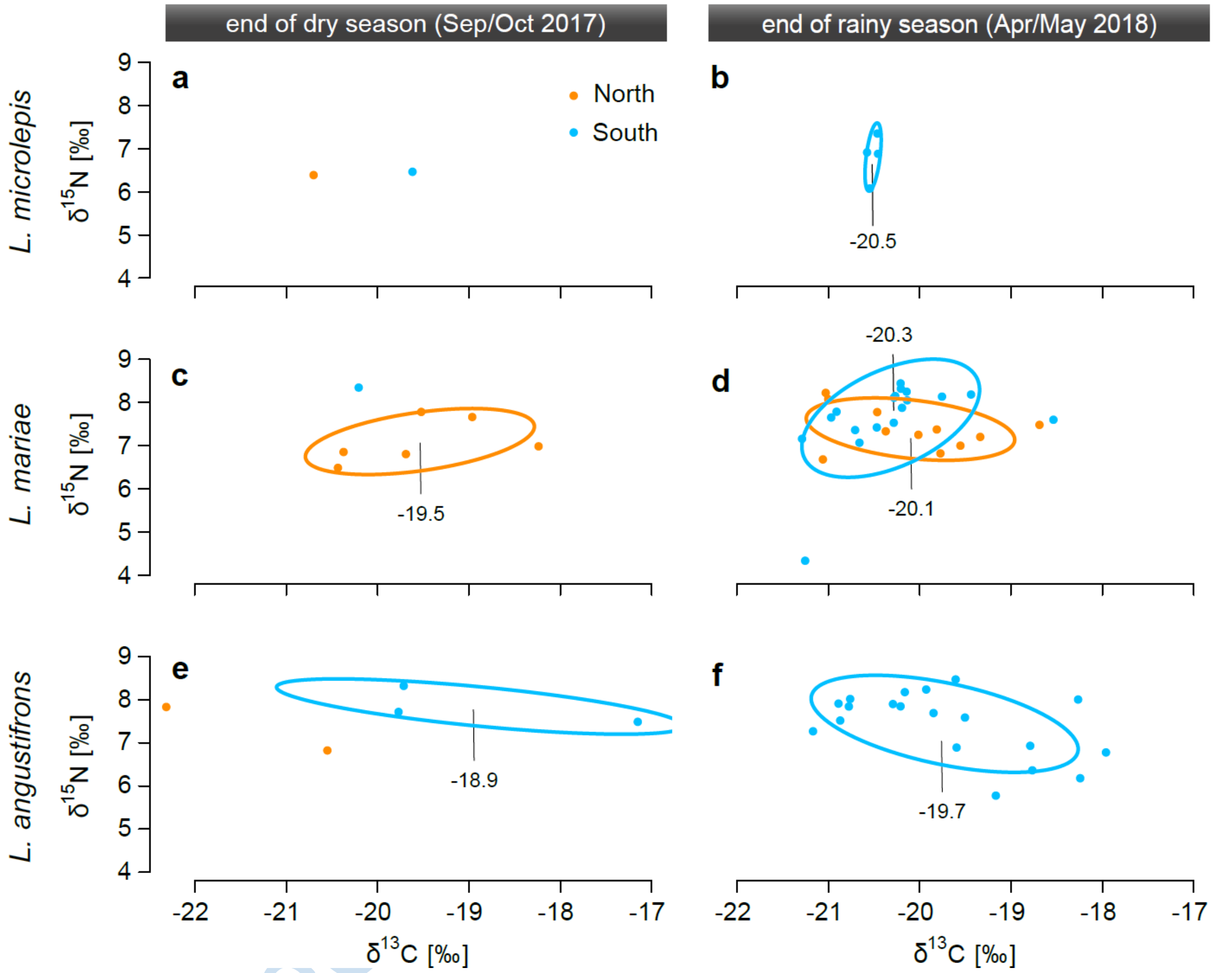


**Fig. S5:** Carbon (normalized for C:N mass ratio according to Post *et al.*, 2007) and nitrogen stable isotope signatures of the large *Lates* species, namely (**a,b**) *Lates microlepis* (**c,d**) *Lates mariae*, and (**e,f**) *Lates angustifrons* at the end of the dry season (left) and the end of the rainy season (right). Orange dots represent the northern basin (stations 1-3) and blue dots represent the southern basin (stations 7-9). Numbers indicate the mean $\delta^{13}C$ of a population. Only individuals >150 mm were included in this analysis to reduce ontogenetic effects on the isotope signatures.

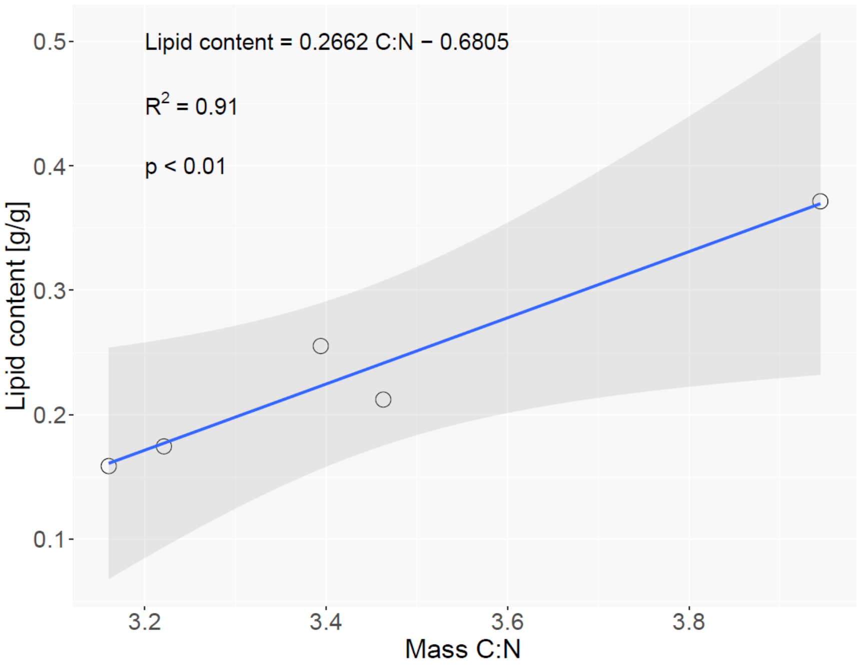


**Fig. S6:** Lipid content versus C:N ratios of *Stolothrissa tanganicae*. Each dot represents a tissue sample from one specimen.

not peer-reviewed preprint

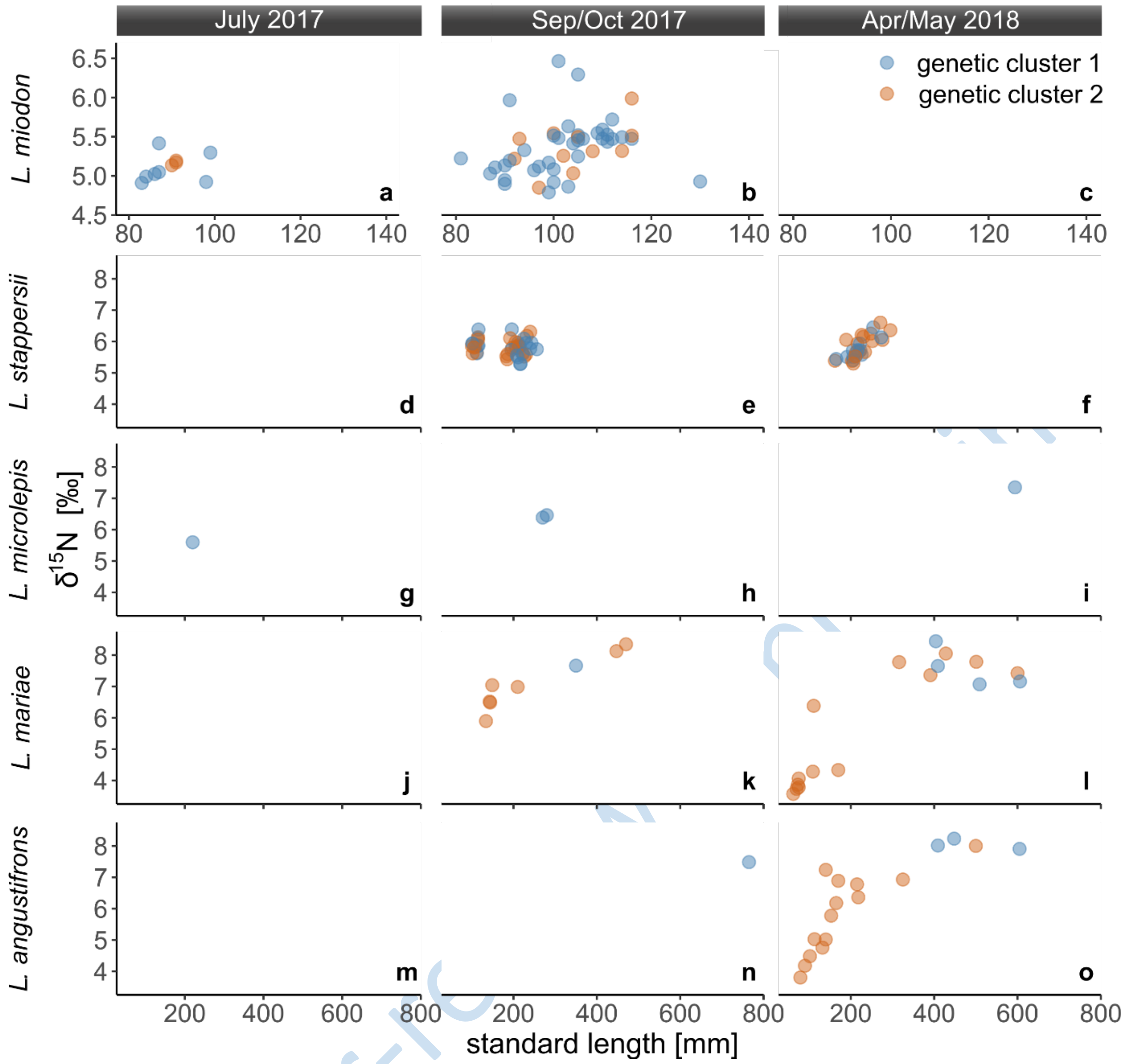


**Fig. S7:** Nitrogen stable isotope signatures of different genetic clusters within the studied fish species in (**left**) July 2017, (**middle**) September/October 2017, and (**right**) April/May 2018. (**a-c**) *Limnothrissa miodon*, (**d-f**) *Lates stappersii*, (**g-i**) *Lates microlepis* (**j-l**), *Lates mariae*, and (**m-o**) *Lates angustifrons*. Note the different axis scaling between *Limnothrissa* and the *Lates* species.

not peer-reviewed
